# Therapy-associated remodeling of pancreatic cancer revealed by single-cell spatial transcriptomics and optimal transport analysis

**DOI:** 10.1101/2023.06.28.546848

**Authors:** Carina Shiau, Jingyi Cao, Mark T. Gregory, Dennis Gong, Xunqin Yin, Jae-Won Cho, Peter L. Wang, Jennifer Su, Steven Wang, Jason W. Reeves, Tae Kyung Kim, Youngmi Kim, Jimmy A. Guo, Nicole A. Lester, Nathan Schurman, Jamie L. Barth, Ralph Weissleder, Tyler Jacks, Motaz Qadan, Theodore S. Hong, Jennifer Y. Wo, Hannah Roberts, Joseph M. Beechem, Carlos Fernandez-del Castillo, Mari Mino-Kenudson, David T. Ting, Martin Hemberg, William L. Hwang

## Abstract

In combination with cell intrinsic properties, interactions in the tumor microenvironment modulate therapeutic response. We leveraged high-plex single-cell spatial transcriptomics to dissect the remodeling of multicellular neighborhoods and cell–cell interactions in human pancreatic cancer associated with specific malignant subtypes and neoadjuvant chemotherapy/radiotherapy. We developed Spatially Constrained Optimal Transport Interaction Analysis (SCOTIA), an optimal transport model with a cost function that includes both spatial distance and ligand–receptor gene expression. Our results uncovered a marked change in ligand–receptor interactions between cancer-associated fibroblasts and malignant cells in response to treatment, which was supported by orthogonal datasets, including an *ex vivo* tumoroid co-culture system. Overall, this study demonstrates that characterization of the tumor microenvironment using high-plex single-cell spatial transcriptomics allows for identification of molecular interactions that may play a role in the emergence of chemoresistance and establishes a translational spatial biology paradigm that can be broadly applied to other malignancies, diseases, and treatments.

## MAIN

Pancreatic ductal adenocarcinoma (PDAC) remains one of the deadliest malignancies with a five-year overall survival of approximately 11%.^1^ Even among patients with locoregional disease, standard of care neoadjuvant chemotherapy with or without radiotherapy followed by surgical resection yields modest outcomes.^2^ Indeed, treatment failure is pervasive, thought to be driven largely by genetic and phenotypic heterogeneity as well as a highly desmoplastic and immunosuppressive microenvironment.^3,4^ Since most patients enrolling in clinical trials have received and progressed through standard of care cytotoxic treatment, identification of novel therapeutic targets in the remodeled tumor microenvironment (TME) would benefit from a deeper understanding of cell–cell interactions among distinct cell types.

Recent applications of single-cell and spatial transcriptomic technologies to PDAC have deepened our understanding of intratumoral heterogeneity, cell state plasticity, therapeutic sensitivities, and neighborhood composition based on co-localization of malignant subtypes with specific stromal and immune subpopulations.^5–12^ However, prior studies were unable to provide high-plex molecular information while preserving *in situ* spatial relationships at single-cell resolution, limiting detailed analyses of cell–cell interactions.^6,7,12–15^ Cell–cell interactions orchestrate the development and function of tissues in normal physiology and disease. Signaling among diverse cell types is often mediated by ligand–receptor (LR) protein interactions involving both soluble and membrane-bound factors. In conjunction with cell intrinsic properties, cell–cell interactions influence many cellular decisions ranging from lineage differentiation to programmed cell death.^16,17^ Cell–cell interactions are also known to play an important, yet incompletely understood, role in the TME, and we hypothesize that changes due to treatment will be partially reflected by changes in LR interactions.

Several methods have been developed to infer cell–cell interactions from single-cell RNA-sequencing (scRNA-seq) data.^17,18^ However, most of these models ignore intercellular proximity *in situ* because scRNA-seq data lacks the spatial information necessary for determining the positions of both sending and receiving cells. Moreover, methods that use scRNA-seq to interrogate physically-interacting cells are unable to capture proximal cells that interact via soluble factors rather than direct physical contact.^15^ Recent advances in spatial transcriptomics have enabled subcellular resolution while profiling up to ∼1,000 transcripts,^19,20^ offering an unprecedented ability to understand both the micro-organization and cell–cell interactions that underpin the pre- and post-treatment TME in their *in situ* context.

To harness the latest advances in spatial transcriptomic technologies, we developed a translational spatial biology paradigm that integrates high-plex and single-cell resolution spatial transcriptomics with specialized computational methods and preclinical models to explore the remodeled TME in response to standard-of-care therapies. We performed a detailed dissection of the cell–cell interactions in the TME associated with neoadjuvant cytotoxic therapy by performing spatial molecular imaging (SMI^19^) on formalin-fixed paraffin-embedded (FFPE) treated and untreated primary resected human PDAC. Leveraging the single-cell spatial resolution of SMI, we developed Spatially Constrained Optimal Transport Interaction Analysis (SCOTIA), an optimal transport model with a cost function that considers both spatial distance and LR gene expression. Our analysis highlighted cell-cell interactions associated with specific malignant cell states as well as striking differences in LR interactions between cancer-associated fibroblasts (CAFs) and malignant cells in the treated and untreated settings. We derived additional support for these findings through multiple complementary datasets, including single-nucleus RNA-seq (snRNA-seq) and digital spatial profiling (DSP) of matched samples and an *ex vivo* tumoroid co-culture system. Overall, this study demonstrated that characterization of the TME using high-plex single-cell spatial transcriptomics enables identification of molecular interactions that may play a role in the emergence of chemoresistance.

## RESULTS

### Spatial molecular imaging captures pancreatic cancer transcriptomics at single-cell resolution

We dissected the spatial transcriptomic landscape of primary resected human PDAC tumors with (*n* = 6) or without (*n* = 7) neoadjuvant chemotherapy and radiotherapy (CRT) at single-cell resolution using spatial molecular imaging (SMI) (**Fig. S1A**) and a 960-plex gene panel, with (*n* = 6) and without (*n* = 9) 30 custom probes (**Fig. 1A; Fig. S1B; Table S1; Methods**). Two tumors were profiled by both the base 960-plex and the augmented 990-plex panels on consecutive sections. Custom probes were selected to improve annotation of malignant subtypes previously identified in PDAC (**Methods**).^6^ Neoadjuvant-treated tumors received multi-cycle FOLFIRINOX chemotherapy followed by multi-fraction radiotherapy with concurrent fluorouracil/capecitabine, with (*n* = 1) or without losartan (*n* = 5). Of the 13 independent tumors in this study, matched single-nucleus RNA-sequencing (snRNA-seq) and digital spatial profiling (DSP; Nanostring GeoMx) data was available for 8 and 10 samples, respectively (**Fig. 1A**).^6^

**Fig. 1.**
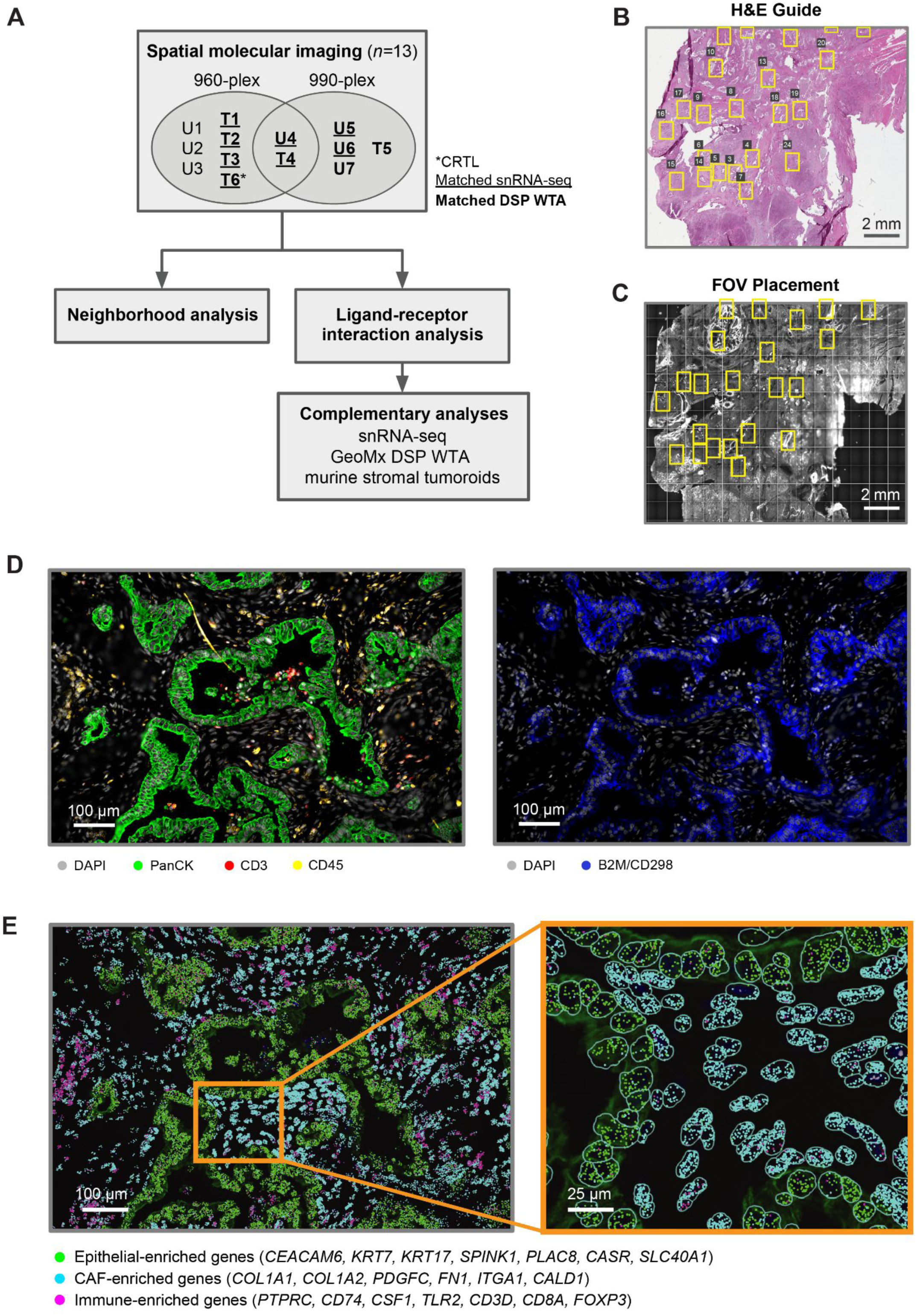
Spatial molecular imaging captures pancreatic tumor architecture at subcellular resolution. **(A)** Study design for human PDAC tumors (*n* = 13) using SMI, followed by computational analyses and experimental investigation of candidate ligand–receptor pairs. Tumors were divided into treatment-naive (*n* = 7) and neoadjuvant-treated (chemotherapy and radiotherapy, *n* = 6) cohorts, with one treated sample additionally receiving losartan. Samples were analyzed using a base 960-plex probe set, with (*n* = 6) and/or without (*n* = 9) 30 custom probes. Two tumors, one treated and one untreated, were profiled using both the base 960-plex and augmented 990-plex panels on consecutive sections. A subset of samples has matched snRNA-seq and whole transcriptome DSP (NanoString GeoMx) data. U, untreated; T, treated. **(B-C)** Representative hematoxylin and eosin (H&E)-stained FFPE section (5 µm thickness) **(B)** and SMI slide image **(C)** of consecutive tissue sections, showing selected fields of view (FOVs, yellow rectangles). The H&E-stained FFPE section was used as a guide for SMI FOV placement to enrich for tumor areas. **(D)** Immunofluorescence image of a representative FOV with antibody targets shown in the color legend. **(E)** Spatial coordinates of RNA transcripts for canonical epithelial (green), CAF (cyan) and immune (magenta) marker genes, overlaid on the immunofluorescence image. *Inset* depicts a magnified view of a region within the FOV, with cell segmentation boundaries (cyan) outlined.

We examined hematoxylin and eosin (H&E)-stained formalin-fixed paraffin-embedded (FFPE) sections (**Fig. 1B**) to guide selection of fields of view (FOVs; 984.96 µm x 662.04 µm) for SMI on consecutive tissue sections (**Fig. 1C**). We spatially localized proteins (antibodies against pan-cytokeratin, CD3, CD45, CD298/B2M, and DAPI) (**Fig. 1D**) and messenger RNA transcripts (**Fig. 1E**) across 320 FOVs. We applied a morphology-based cell segmentation algorithm^19^ to define cell boundaries (**Fig. 1E** inset), and removed cells with fewer than 60 total transcripts (**Methods**). This yielded 753,133 high confidence single-cell profiles.

To characterize the spatial organization of different cell types in PDAC, we first annotated cells using the Insitutype supervised clustering method^21^ and snRNA-seq data from our previous study as a reference^6^ (**Fig. 2A**; **Methods**). More than 95% of cells were successfully annotated to a distinct cell type, and the diversity of cell types in PDAC was captured, including malignant, cancer-associated fibroblasts (CAFs), endothelial, pericyte, vascular smooth muscle, endocrine, Schwann/nerve, myeloid, and lymphoid cells (**Fig. 2B-D; Fig. S2A-B**). The median number of cells in untreated and treated FOVs was 2,718 and 1,645, respectively, consistent with cell death secondary to cytotoxic treatment. There were significantly more malignant cells in untreated compared to treated tumors (median of untreated FOVs: 1,152, treated FOVs: 269; *p* < 10^−16^, two-sided Mann-Whitney U; **Fig. S2C**), but there was no significant difference across treatment groups for non-malignant cells (median of untreated FOVs: 1,476, treated FOVs: 1,245; *p* = 2.18×10^−1^; **Fig. S2C**). Within non-malignant cells, only CAFs had significantly lower cell proportions in untreated tumors compared to treated tumors (median of untreated FOVs: 0.559, treated FOVs: 0.652; *p* = 1.70×10^−8^, adjusted two-sided Mann-Whitney U; **Fig. S2D**), supporting previous reports that cytotoxic therapies may induce CAF proliferation and remodeling as well as augment the underlying desmoplastic reaction.^6,12,22^ After further sub-clustering of CAFs and malignant cells, we identified five subsets: inflammatory CAFs (iCAF), myofibroblastic CAFs (myCAF), classical (CLS) malignant cells, basal-like (BSL) malignant cells, and the recently characterized neural-like progenitor (NRP) malignant subtype that exhibits stem-like and neurodevelopmental features (**Fig. 2B, E-G**).^6,23–25^ The aggregate basal-like subtype applied in this analysis could not be confidently resolved into distinct basaloid, squamoid, and mesenchymal states^6^ due to limitations in the panel size. Consistent with prior work,^6^ CLS and BSL were the two most common malignant subtypes, with NRP cells only constituting 6.23% of all malignant cells.

**Fig. 2.**
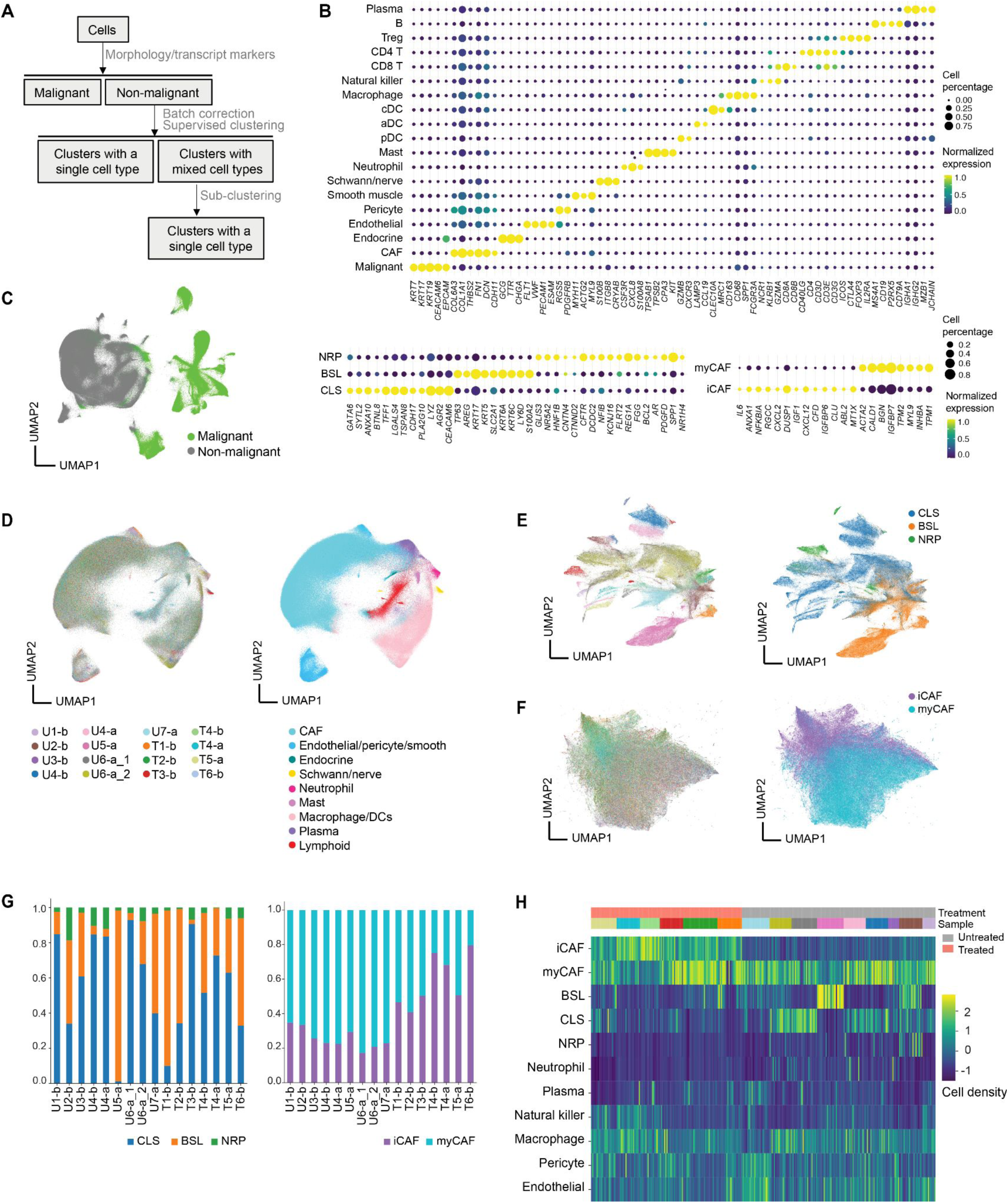
Spatial molecular imaging uncovers cell type diversity in pancreatic cancer. **(A)** Schematic of supervised cell typing procedure (**Methods**). **(B)** Bubble heatmap showing expression levels of select marker genes for annotated cell types and subtypes. Color indicates normalized expression and dot size indicates the fraction of expressing cells. Treg, regulatory T cell; cDC, conventional dendritic cells; aDC, activated dendritic cells; pDC, plasmacytoid dendritic cells; CAF, cancer-associated fibroblast; NRP, neural-like progenitor; BSL, basal-like; CLS, classical; myCAF, myofibroblastic CAF; iCAF, inflammatory CAF. **(C)** UMAP visualization of malignant and non-malignant cells. **(D)** UMAP visualization of batch-corrected non-malignant cells colored by sample ID (*left*) and cell type annotation (*right*). U, untreated; T, treated; b, base 960-plex panel; a, augmented 990-plex panel. Sample ID color legend shared with panels E and F. **(E)** UMAP showing malignant subsets colored by patient ID (*left*) and subtype annotations (*right*). **(F)** UMAP showing CAF subsets colored by sample ID (*left*) and subtype annotations (*right*). **(G)** Proportions of malignant (*left*) and CAF (*right*) subtypes across untreated and treated tumors. **(H)** Heatmap showing Z score normalized cell type densities (color bar, *right*) for each FOV across treatment status (color bar labeled “Treatment,” *top*) and specimens (color bar labeled “Sample”, *top*).

We next asked whether cell type composition is predictive of whether a FOV was from an untreated or treated sample. Using the cell type annotations (**Fig. 2C-D**), we trained a *k* nearest neighbor classifier that used the densities of 11 cell types or subtypes (**Fig. 2H**). Holding out all FOVs from one of the fifteen samples during training, an accuracy of 0.79 was achieved. Closer inspection of the distribution of cell types between the two conditions revealed that although neutrophils represent only 0.56% of cells, 92.5% were found in untreated samples. Concordantly, we observed significantly lower expression of chemokines associated with neutrophil chemotaxis (*CXCL1/2/3/5/6/8*) in both malignant cells (*log_2_*FC = −1.79, *p* < 10^−16^, two-sided Mann-Whitney U; **Fig. 2E**) and CAFs (*log_2_*FC = −0.54, *p* = 2.60×10^−3^; **Fig. S2E**) in treated samples, indicating that the lower abundance of neutrophils in treated samples may be partially attributed to reduced recruitment.^26^ Circulating neutrophil counts in the peripheral blood do not significantly differ (*p* = 0.282, Mann-Whitney U) between treated (2977±1300 neutrophils/µL) and untreated (4974±3338 neutrophils/µL) patients, further supporting the interpretation that the lower neutrophil density in treated samples is due to reduced neutrophil recruitment into the TME rather than systemic neutrophil depletion.

While cell density is a global metric capturing the overall heterogeneity among FOVs, it does not take advantage of spatial positions. To leverage this spatial information, we developed a novel feature extraction strategy based on topological data analysis (**Methods**).^27^ Briefly, this approach characterizes the length scales separating each different cell type and uses persistence images^28^ to convert this information into a metric space suitable for machine learning. For each cell type we obtained a one-dimensional persistence image with five features, and the images were combined for a *k* nearest neighbors classifier. Using the same training and testing scheme as before, this classifier was able to distinguish FOVs based on treatment with an accuracy of 0.84. This result demonstrates that it is not only the cell density, but also the spatial distribution of cells that differs between the treated and untreated samples. Together, these results support the conclusion that our annotation pipeline captures the complexity and heterogeneity of PDAC.

### Neighborhood analyses reveal low glandular heterogeneity and specific multicellular associations

The previous analysis indicated a quantitative difference in overall tissue architecture as a function of treatment, so we further investigated the subtype composition of malignant glands. To extract individual glands, we separately applied DBSCAN^29,30^ to the pan-cytokeratin antibody channel and the spatial locations of malignant cells to obtain two independent clustering assignments. We acquired a consensus between the results by merging distinct clusters in one assignment that have at least one cell in the same cluster by the other assignment method (**Fig. 3A; Fig. S3A**; **Methods**). As expected, there were significantly fewer glands in the treated samples (*p* < 10^−16^, two-sided Mann-Whitney U) (**Fig. 3B-C**). We then compared the number of malignant cells and the proportion of malignant subtypes across glands (**Fig. 3B-C**). Glands in treated samples were composed of fewer cells than those in untreated samples (*p* = 3.47×10^−5^, two-sample K-S), while treated samples had a significantly greater proportion of singular malignant cells than untreated samples (*p* = 2.04×10^−12^, two-sample K-S), suggesting that malignant cells are less inclined to aggregate into glandular structures after treatment (**Fig. 3C**). Most glands were predominantly composed of a single malignant subtype (65.9% of glands), while others were composed of a mixture of subtypes (**Fig. 3B**; **Methods**). CLS-predominant glands were significantly larger than BSL-predominant glands (*log_2_*FC = 1.427; *p* < 10^−16^, two-sample K-S) and NRP-predominant glands (*log_2_*FC = 1.435; *p* = 1.05×10^−13^, two-sample K-S) (**Fig. 3D**; **Methods**). Since most glands were dominated by one subtype, the proportion of CLS and BSL cells in each gland was significantly anti-correlated (Spearman *r* = −0.678, *p* < 10^−16^).

**Fig. 3.**
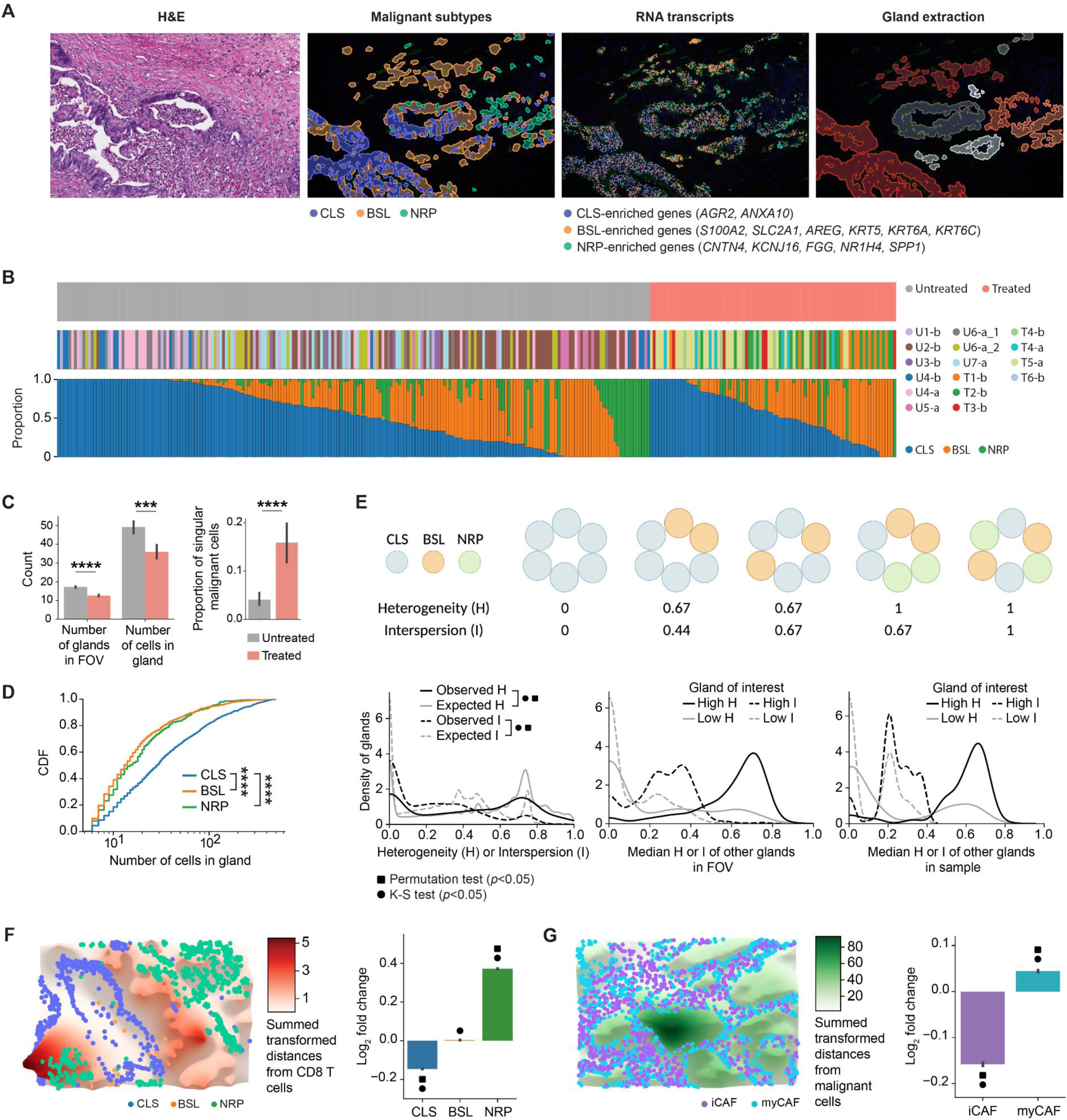
Spatial molecular imaging reveals glandular heterogeneity and multicellular neighborhoods in pancreatic cancer. **(A)** (*Left to right*) Hematoxylin and eosin (H&E) stain, SMI-identified malignant cells (colored by subtype), localized RNA transcripts for a subset of malignant subtype marker genes, and DBSCAN-identified malignant glands (colored in shades of red and gray) for a representative FOV. **(B)** Treatment condition (color bar, *top*), sample ID (color bar, *middle*) and malignant subtype proportion (stacked proportion bar plot, *bottom*) for a random subset of malignant glands (*n*=300, 10% of total glands). U, untreated; T, treated; b, base 960-plex panel; a, augmented 990-plex panel. **(C)** Number of glands per FOV (*left)*, number of cells per gland (*middle*), and the proportion of singular malignant cells (*right*), stratified by treatment condition. Error bars denote the 95% confidence interval. \*\*\**p* < 0.001; \*\*\*\**p* < 0.0001, two-sample Mann-Whitney U test. **(D)** Cumulative density function for the number of cells in malignant glands, separated malignant subtype. \*\*\*\**p* < 0.0001, two-sample K-S test. **(E)** *Top*: Schematic of the subtype composition of representative malignant glands, with heterogeneity and interspersion measurements noted (**Methods**). *Bottom*: Observed (black) and expected (gray) distributions of the heterogeneity (H, solid line)/interspersion (I, dotted line) of malignant glands (*left*) or the median H/I of all other glands in the FOV (*middle*) and sample (*right*) for glands in the top *versus* bottom quartile of H/I. Statistical test legend shared with panels F and G. **(F-G)**, *Left*: Three-dimensional topological depiction of the summed exponential functions (*z* axis height and color bar, decay radius *r* = 50 μm) that are generated from CD8 T **(F)** and malignant **(G)** cells for a representative FOV, with spatial locations of malignant **(F)** and CAF **(G)** subtypes shown as colored dots. *Right*: Log_2_ fold change (*y* axis; error bars denote the 95% confidence interval) between the observed and expected summed exponential-transformed distances between malignant subtypes and CD8 T cells (using a malignant-centric model) **(F)**, and between CAF subtypes and malignant cells (using a CAF-centric model) **(G)**.

Interestingly, the composition of malignant glands was more homogeneous than expected by chance (*p <* 10^−16^, two-sample K-S; *p* < 0.001, permutation test) (**Fig. 3E**), with glands from untreated samples being more homogeneous than those from treated samples (*p* = 2.32×10^−14^, two-sample K-S) (**Methods**). Moreover, heterogeneous glands were more likely to be found in the same FOV as other heterogeneous glands, while homogeneous glands tended to co-localize with homogeneous glands (**Fig. 3E**). In addition to quantifying heterogeneity, we measured gland interspersion (**Methods**), which provides information about the degree of subtype mixing within a gland. Malignant glands were less interspersed than expected by chance (*p* < 10^−16^, two-sample K-S; *p* < 0.001, permutation test), and we similarly observed that highly interspersed glands tended to co-localize with other highly interspersed glands, and vice versa. These findings were consistent at both the FOV and specimen level (**Fig. 3E**). Moreover, the organization of malignant subtypes within highly heterogeneous glands was significantly less interspersed than expected (*p <* 10^−16^, two-sample K-S; p < 0.001, permutation test) (**Fig. S3B**).

Next, we developed a model based on an exponential to measure the pairwise proximity of different cell types (**Fig. 3F-G**; **Methods**). We applied this model to investigate a prior low-resolution spatial association between NRP malignant cells and CD8 T cells.^6^ To quantify the spatial relationship between CD8 T cells with each malignant subtype at single-cell resolution, we measured the summed exponential-transformed distances between CD8 T cells and malignant cells (**Fig. 3F; Methods**). We compared the observed spatial distributions to a null distribution obtained by shuffling the malignant annotations within each FOV. This analysis revealed that CD8 T cells were closer to NRP malignant cells than expected by chance (*log_2_*FC = 0.371±0.0200; *p* < 10^−16^, two-sample K-S; *p* < 0.001, permutation test), while CD8 T cells were located further from CLS malignant cells than expected (*log_2_*FC = −0.146±0.00551; *p* = 1.93×10^−7^, two-sample K-S; *p* < 0.001, permutation test) (**Fig. 3F; Fig. S3C**), which was concordant with our prior findings.^6^ These associations were robust to the selection of parameters for the exponential function as well as to the choice of malignant-versus CD8 T cell-centric models (**Fig. S3D-G**; **Methods**). Additionally, we found that myCAFs (*log_2_*FC = 0.0432±0.00669; *p* < 10^−16^, two-sample K-S; *p* < 0.001, permutation test) were more proximal to malignant cells than iCAFs (*log_2_*FC = −0.158±0.0114; *p* < 10^−16^, two-sample K-S; *p* < 0.001, permutation test), recapitulating a prior observation (**Fig. 3G; Fig. S3H**).^24^

### Identification of spatially defined CAF–malignant interactions enriched in treated tumors

We hypothesized that it is not only the cellular architecture, but also the cellular interactions that differ between treatment conditions. To understand how ligand–receptor (LR) interactions vary among cell types and conditions, we developed a computational method that integrates the expression levels of ligands and receptors with spatial distance by optimal transport (OT; **Fig. 4A**; **Methods**). We refer to the method as Spatially Constrained Optimal Transport Interaction Analysis (SCOTIA), which is freely available (see “Data and Code Availability” section in **Methods**). SCOTIA first extracts spatial clusters of cells by type, and then excludes cluster pairs that are far apart and hence less likely to interact. The cell–cell interaction strength is inferred by minimizing the total transport cost between the adjacent source and target cell clusters (**Fig. 4A**). Since the original OT algorithm only considers spatial distance as transport cost, we modified the cost matrix to incorporate distance in gene expression space as well (**Methods**). Although analyses at the level of malignant and CAF subtypes would be interesting, similar ligand-receptor expression among subtypes (**Fig. S4A**) and low prevalence of certain subtypes (**Fig. 2G**) compromises statistical power. Hence, we focused on analyzing interactions among major cell types. Systematic investigation of cataloged LR pairs between any two cell types (**Fig. S4B-C**; **Methods**) revealed that the largest difference in interaction strength was between CAFs and malignant cells when comparing treated residual tumors to untreated tumors (**Fig. 4B-C**). The strongest effect was observed for CAF ligands signaling to malignant cell receptors, and this result was robust to the choice of cost function (**Fig. S4D**). These results suggest a treatment-associated role for CAFs in modulating malignant cells and the tumor microenvironment.^13,31,32^

**Fig. 4.**
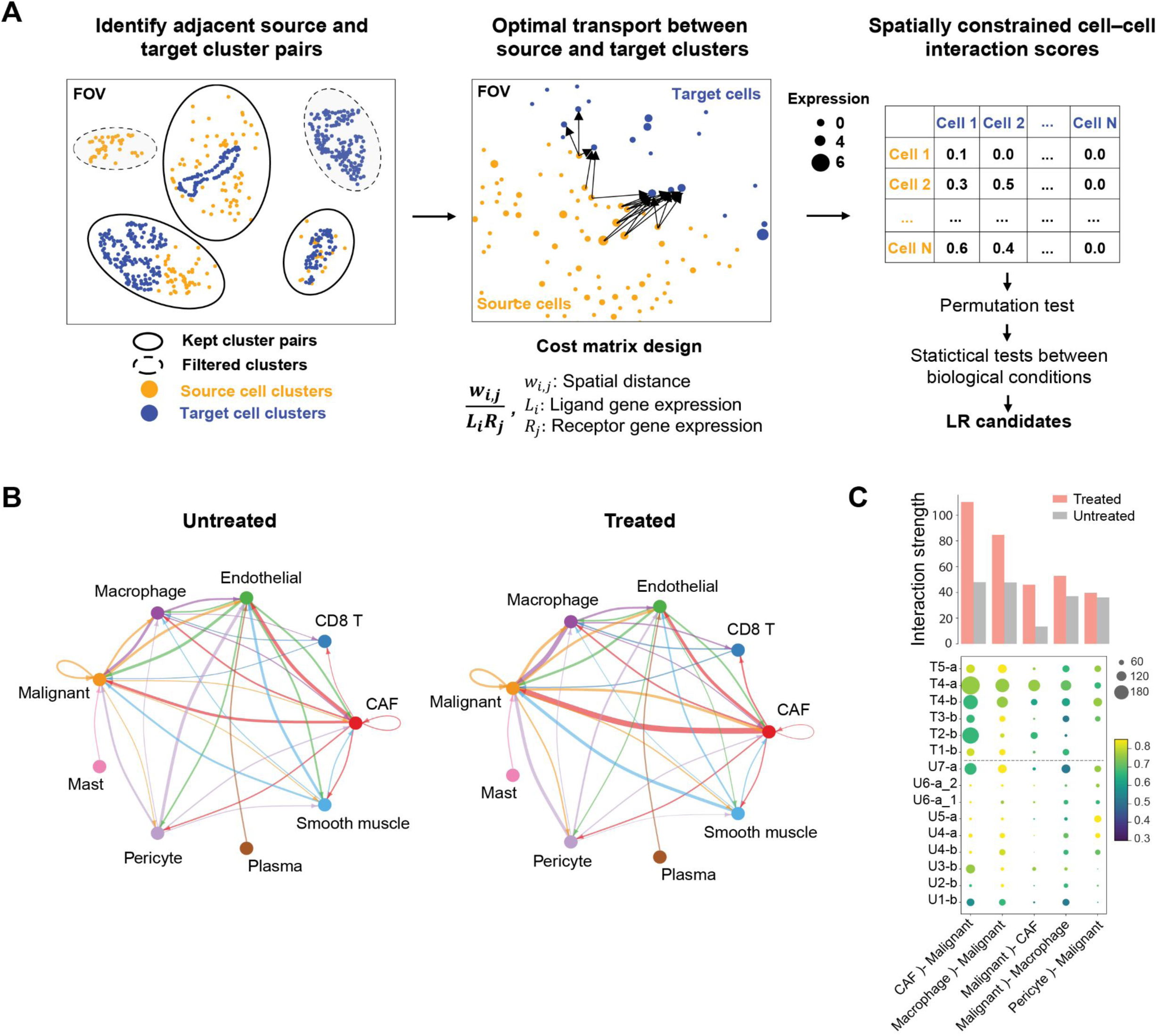
Deciphering cell–cell interactions using Spatially Constrained Optimal Transport Interaction Analysis (SCOTIA). **(A)** Overview of the workflow for inferring ligand–receptor (LR) interactions from spatial molecular imaging data (**Methods**). Cells were clustered using DBSCAN for each cell type and only spatially adjacent source and target cell clusters were retained for downstream LR analysis (*left*). LR interaction scores were calculated using an optimal transport (OT) model in which the cost matrix integrates spatial distance and LR expression (*middle*). Statistical tests were performed to identify treatment-associated LR candidates (*right*). **(B)** The inferred total interaction strength between annotated cell type pairs for untreated (*left*) and treated (*right*) tumors. Interaction strength was measured as the summed LR interaction scores for permutation test-significant LR pairs. Edge width is proportional to interaction strength and edge color reflects source cell type. **(C)** The top five strongest interacting cell type pairs from panel **(B)** are shown for each tumor. Dot size represents the number of permutation test-significant LR pairs, colored based on the average LR interaction score. Bar plot indicates the average interaction strengths of each cell type pair for the treated and untreated groups. U, untreated; T, treated; b, base 960-plex panel; a, augmented 990-plex panel.

### Cytotoxic treatment alters ligand–receptor interactions between CAFs and malignant cells

To explore whether neoadjuvant therapy was associated with significant differences in interactions between CAF ligands and malignant receptors, we identified 22 LR pairs that were significantly enriched and 54 LR pairs that were significantly depleted in treated tumors (adjusted *p* < 0.05, two-sided Mann-Whitney U; **Fig. 5A-B; Fig. S5A-B**). Visual inspection confirmed the inferred differences in spatial and gene expression distance between LR pairs as a function of treatment status (**Fig. 5C**). Treatment-enriched LR interactions (**Fig. 5B**) were associated with chemotaxis, cytokine signaling, stromal/extracellular matrix (ECM) remodeling, and immune modulation (**Fig. S5C**). Interestingly, several ligands, including TGFB1 and CXCL12, and receptors, including CXCR3, CXCR4, and ACKR3, have been associated with therapeutic resistance.^33–37^ We also identified a treatment-associated enrichment in IL6 family signaling and JAK/STAT activation (e.g., *CLCF1–CNTFR, LIF–IL6ST, CSF2–CSF3R*), which has been linked to epithelial-to-mesenchymal transition (EMT), invasion, metastasis, immune modulation, and resistance to chemotherapy and radiotherapy in numerous malignancies.^38–47^

**Fig. 5.**
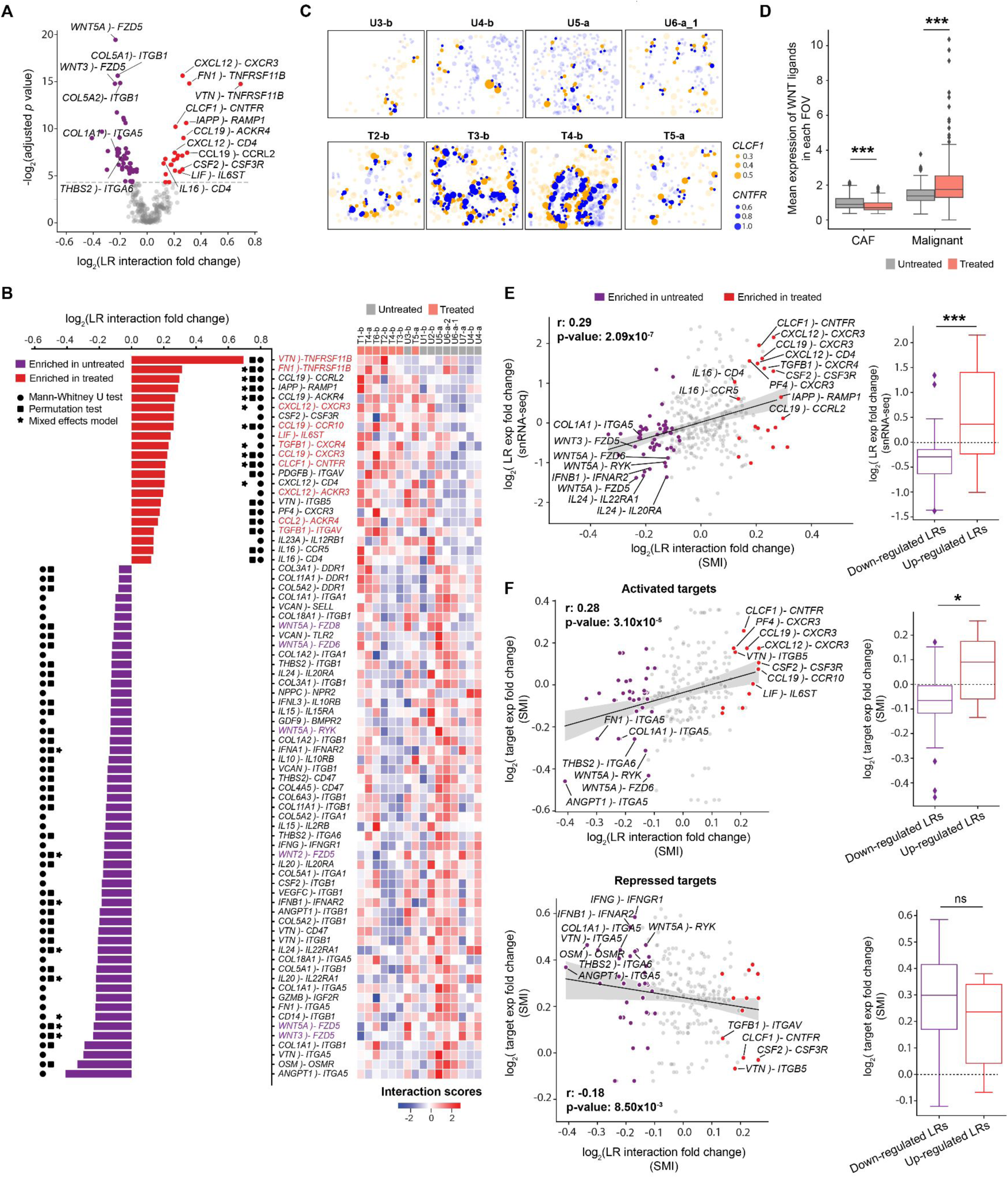
Treatment-associated ligand–receptor interactions between CAFs and malignant cells. **(A)** Volcano plot showing log_2_ fold change of ligand–receptor (LR) interaction scores (*x* axis) between treated and untreated samples versus –log_2_ Benjamini-Hochberg adjusted *p* values calculated from two-sided Mann-Whitney U test (*y* axis). In this analysis, CAFs serve as sources and malignant cells act as targets. LR interactions that are significantly up- or down-regulated in treated samples are highlighted in red or purple, respectively. **(B)** (*Left*) Waterfall plot showing the significant differentially-enriched LR pairs from panel **(A)**. Significant results (adjusted *p* < 0.05) of Mann-Whitney U, permutation, and mixed effects model tests were indicated with circle, square, and asterisk symbols, respectively. (*Right*) Heatmap showing the hierarchical clustering of patients with z-score normalized LR interaction scores. U = untreated; T = treated; b = base 960-plex panel; a = augmented 990-plex panel. **(C)** Spatial visualization of an example treatment-enriched LR interaction in representative FOVs: *CLCF1–CNTFR*. Dots represent ligand gene expression in CAF cells (orange) or receptor gene expression in malignant cells (blue) with the size indicating normalized expression levels and the locations indicating spatial coordinates of cell centroids. Cells expressing ligands or receptors but not participating in CAF–malignant interactions are shown as semi-transparent dots. **(D)** Expression of WNT ligands in treated/untreated CAFs and malignant cells. WNT ligands include *WNT2/2B/3/5A/5B/7A/7B/9A/10B/11*. Two-sided Mann-Whitney U test (\*\*\**p* < 0.001). **(E)** (*Left*) Scatter plot showing the correlation between the log_2_ fold change of LR interaction scores from SMI data (*x* axis) and the log_2_ fold change of LR gene expression in the published snRNA-seq dataset^6^ (*y* axis). LR pairs were highlighted with the same color scheme as panel **(A)**. LR gene expression in the snRNA-seq data was calculated by averaging ligand expression in CAFs and receptor expression in malignant cells. Spearman rho and *p* values were provided. (*Right*) Boxplot summarizing the difference in snRNA-seq target gene expression^6^ between significantly enriched and depleted LR pairs identified by applying SCOTIA to the SMI data. Two-sided Mann-Whitney U test (\**p* < 0.05; \*\*\**p* < 0.001; ns, not significant). **(F)** Scatter plots showing the correlations between log_2_ fold change of LR interaction scores (*x* axis) and the log_2_ fold change of downstream target gene expression in the SMI data. *Top*: activated targets; *Bottom*: repressed targets. Spearman rho and *p* values were provided.

On the other hand, systematic analysis of the depleted LR pairs revealed that one of the most prominently represented groups (31 out of 54 pairs) corresponded to integrin-mediated responses in malignant cells stimulated by collagen, angiogenic factors, or hematopoietic growth factors from CAFs (**Fig. 5A-B; Fig. S5C**). Another depleted group of interactions (6 out of 54) was composed of WNT signaling factors (i.e., *WNT3A–FZD5, WNT5A–FZD5, WNT2–FZD5, WNT5A–RYK, WNT5A–FZD6, WNT5A–FZD8*) (**Fig. 5A-B**). We also found that treated malignant cells expressed significantly higher levels of WNT ligands compared to untreated malignant cells (*log_2_*FC = 0.61, *p* = 1.08×10^−5^, two-sided Mann-Whitney U), treated CAFs (*log_2_*FC = 1.43, *p* < 10^−16^), and untreated CAFs (*log_2_*FC = 1.14, *p* < 10^−16^) (**Fig. 5D**). Also, the expression of WNT ligands was lower in treated CAFs compared to untreated CAFs (*log_2_*FC = −0.30, *p* = 2.48×10^−6^; **Fig. 5D**). Several interferon signaling pairs (e.g., *IFNA1–IFNAR2, IFNB1–IFNAR2*, *IFNG–IFNGR1*), and cytokine interactions (e.g., *IL15–IL15RA, IL10–IL10RB, IL24–IL22RA1*, and *IL20–IL20RA*), were also attenuated after treatment. These results suggest that CAF–malignant cell interactions in untreated tumors exhibit high levels of paracrine Wnt signaling, ECM-integrin crosstalk, interferon signaling, and specific subsets of cytokine signaling that may be disrupted after chemotherapy and radiotherapy. Taken together, we have demonstrated that SCOTIA integrates single-cell spatial and gene expression information to identify LR candidates altered in response to therapy.

### Ligand–receptor interactions supported by complementary analyses and co-culture model

To further substantiate the predicted treatment-modulated interactions, we first examined the expression levels of these LR pairs in a prior snRNA-seq dataset.^6^ Reassuringly, the fold change in expression levels was significantly correlated (Spearman *r* = 0.29, *p* = 2.09×10^−7^; two-sided Fisher exact test: *p* = 1.34×10^−4^; two-sided Mann-Whitney U: *p* = 7.06×10^−4^), and several of the most differential interactions, including *CLCF1–CNTFR* and *WNT5A–FZD5*, showed consistent profiles across these datasets (**Fig. 5E**). Next, we investigated the correlation between receptor gene expression and the expression levels of their downstream targets as predicted by the KEGG^48^ and TRRUST^49^ databases (**Methods**). We observed positive and negative correlations for activated (Spearman *r* = 0.28, *p* = 3.10×10^−5^; two-sided Fisher exact test: *p* = 1.65×10^−2^; two-sided Mann-Whitney U: *p* = 2.05×10^−2^) and repressed (Spearman *r* = −0.18, *p* = 8.50×10^−3^; two-sided Fisher exact test: *p* = 1.64×10^−1^; two-sided Mann-Whitney U: *p* = 1.37×10^−1^) target genes (**Fig. 5F**). Additionally, we carried out this downstream target gene analysis for five other abundant cell type pairs and found similar levels of correlation (**Fig. S5D**), suggesting that SCOTIA can reliably infer changes in LR interactions. Furthermore, we ran SCOTIA using two other LR databases, CellPhoneDB^50^ and CellChatDB,^17^ to assess the impact of different inference sources on the identified LR interactions. As expected, the resultant interactions were not entirely overlapping, but nevertheless there was concordance in the biological signaling pathways enriched and depleted with treatment across LR databases (**Table S2**).^51^

Next, we sought to determine whether a subset of the treatment-associated LR pairs identified by SCOTIA would be recapitulated using a murine-derived malignant–CAF co-culture tumoroid system (**Fig. 6A**; **Methods**). We subjected tumoroids to *ex vivo* chemotherapy (5-FU) and separately performed snRNA-seq on treatment-naive and 5-FU treated tumoroids (**Fig. 6A**; **Methods**). We obtained a total of 26,692 high-quality single nucleus profiles (16,200 untreated and 10,492 treated cells), which we clustered into distinct populations of malignant cells and CAFs (**Fig. 6B**). Consistent with the human SMI data analyzed by SCOTIA, LR pairs upregulated in treated co-cultures were enriched in chemotaxis and other cell migration-related features, whereas LR pairs highly expressed in untreated CAFs and malignant cells were related to ECM function and the PI3K pathway (**Fig. S5C, S6A**). Indeed, the fold change of LR expression in treated versus untreated tumoroids was significantly higher for treatment-enriched LR pairs found in the human SMI cohort (*p* = 3.17×10^−2^, two-sided Mann-Whitney U; *p* = 1.01×10^−1^, two-sided Fisher exact test; **Fig. 6C**). Closer inspection revealed that several of the top treatment-enriched LR pairs from the SMI analysis were also upregulated by treatment in the tumoroid snRNA-seq dataset, including *CLCF1–CNTFR*, *CSF2–CSF3R*, and *CCL19–CCRL2*. We also confirmed in both datasets that collagen-integrin interactions, such as *COL1A1–ITGB1, COL1A1–ITGA5*, and *COL3A1–ITGB1*, were dampened in treated samples compared to untreated samples. Overall, these findings demonstrate that the treatment-associated LR pairs we identified using SMI are robust across species and experimental context.

**Fig. 6.**
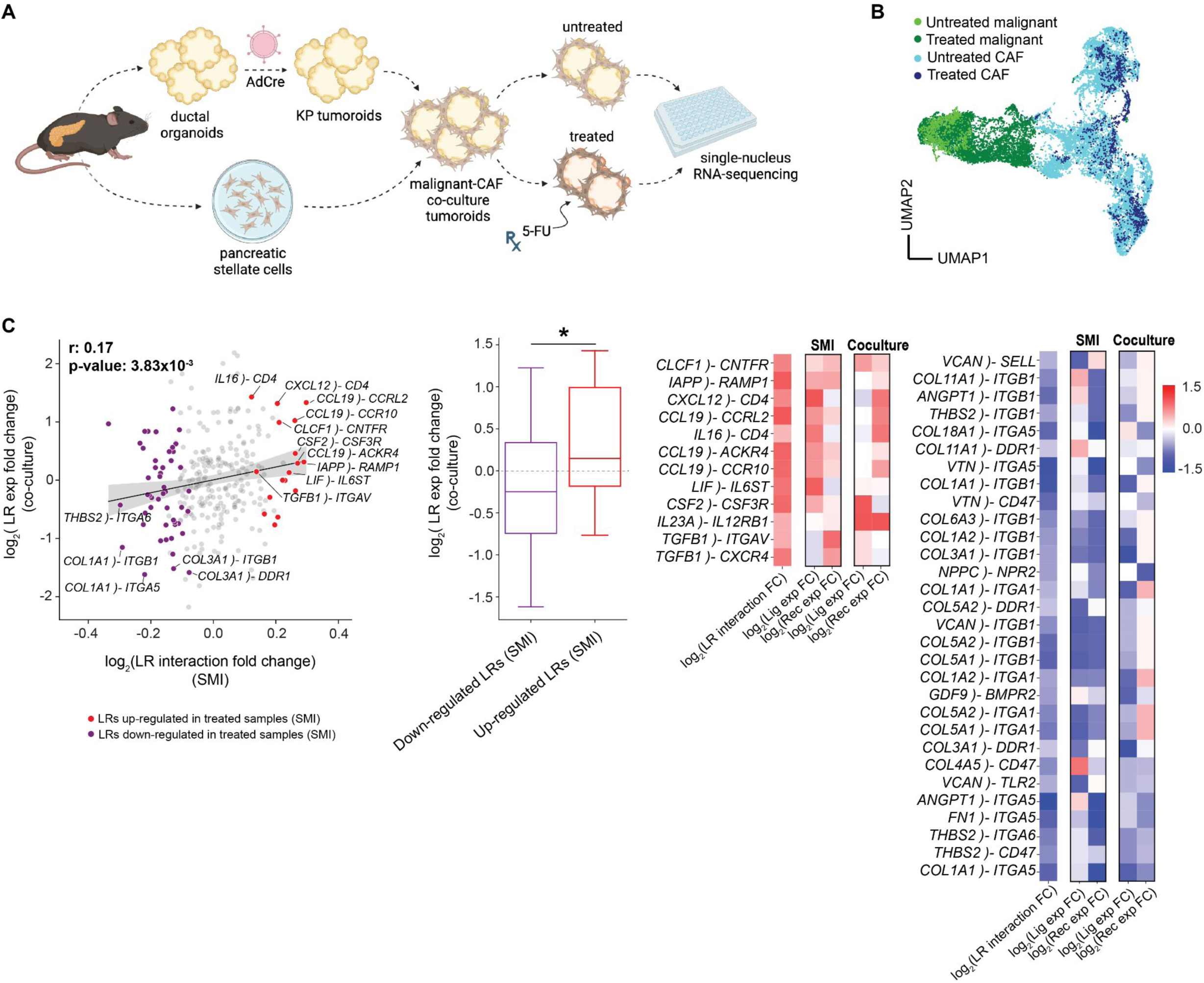
Substantiation of ligand–receptor interactions in murine co-culture tumoroids. **(A)** Experimental workflow for the malignant–CAF co-culture tumoroid system established from KP (*Kras*^G12D/wt^;*Trp53*^FL/FL^) ductal cells and pancreatic stellate cells, both of which were isolated from the mouse pancreas (**Methods**). Single-nucleus RNA-sequencing (snRNA-seq) was performed on untreated and 5-FU-treated malignant-CAF tumoroids. **(B)** UMAP showing batch-corrected snRNA-seq of CAFs and malignant cells dissociated from malignant-CAF tumoroids at steady-state or after treatment with 5-FU. **(C)** (*Left*) Scatter plot showing the correlation between the log_2_ fold change of LR interaction scores in the SMI data (*x* axis) and the log_2_ fold change of LR gene expression in the co-culture dataset (*y* axis). LR interactions that were significantly enriched/depleted in SMI-analyzed treated samples were highlighted in red and purple, respectively. LR gene expression in the co-culture data was calculated by averaging ligand expression in CAFs and receptor expression in malignant cells. Spearman rho and *p* values were provided. (*Middle*) Boxplot summarizing the difference in LR gene expression (*y* axis; co-culture) between the up- and down-regulated LR groups (*x* axis; SMI). Two-sided Mann-Whitney U test (\**p* < 0.05). (*Right*) Heatmap showing a subset of candidate LR interactions that overlap between the SMI and co-culture datasets. Color bar denotes the log_2_ fold change in interaction strength between treated and untreated samples.

### IL-6 family signaling is enriched between inflammatory CAFs and specific malignant subtypes

Among the CAF-malignant interactions enriched in treated samples for both the SMI and co-culture datasets, we observed several members of the IL6 family, including *CLCF1–CNTFR* and *LIF–IL6ST*. Prior studies established a distinction between IL-6^high^ CAFs (iCAFs) and IL-6^low^ CAFs (myCAFs).^24^ Although we observed that iCAFs were further away from malignant cells compared to myCAFs overall (**Fig. 3G; Fig. S3H**), stratification by malignant subtype revealed that NRP malignant cells exhibited the opposite pattern, colocalizing with iCAFs (*log_2_FC* = 0.431 ± 0.0277; *p* < 10^−16^, two-sample K-S; *p* < 0.001, permutation test) more than myCAFs (**Fig. 7A; Fig. S7A**). In addition to NRP malignant cells, we also observed that iCAFs were preferentially localizing with BSL malignant cells, albeit to a lesser degree (*log_2_FC* = 0.0949±0.0102; *p* < 10^−16^, two-sample K-S; *p* < 0.001, permutation test) (**Fig. 7A; Fig. S7A**). iCAFs were significantly enriched in treated compared to untreated samples in both the SMI (*log_2_*FC = 1.10, *p <* 10^−16^, two-sided Mann-Whitney U) and snRNA-seq datasets (*log_2_*FC = 1.28, *p =* 3.80×10^−5^, two-sided Mann-Whitney U) (**Fig. S7B**).^6,12^

**Fig. 7.**
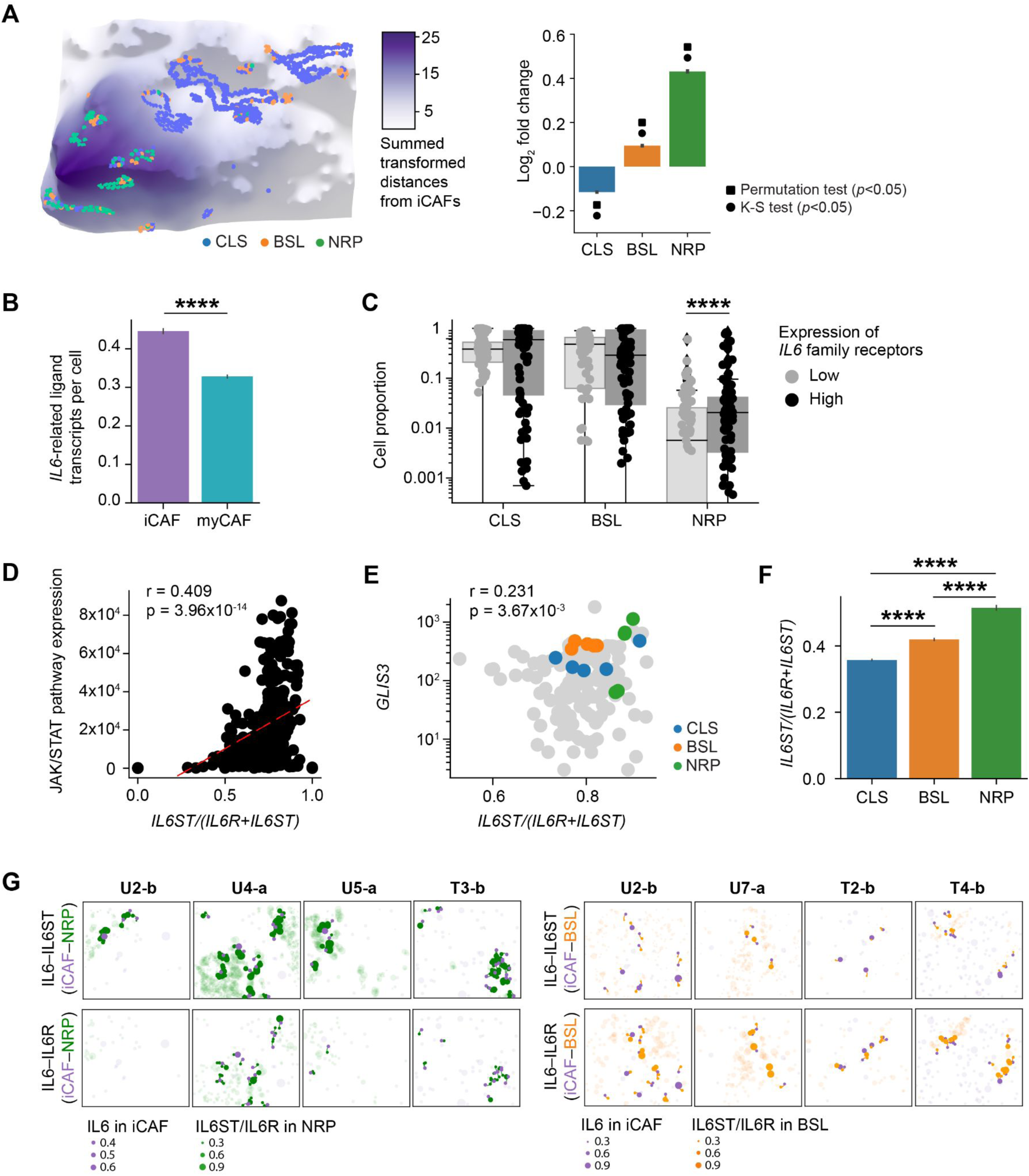
IL-6 family signaling is enriched between inflammatory CAFs and specific malignant subtypes. **(A)** *Left*: Three-dimensional topological depiction of the summed exponential functions (*z* axis height and color bar, decay radius *r* = 50 μm) generated from iCAFs for a representative FOV, with spatial locations of malignant subtypes shown as colored dots. *Right*: Log_2_ fold change (*y* axis; error bars denote the 95% confidence interval) between the observed and expected summed exponential-transformed distances between malignant subtypes and iCAFs. Significant results (adjusted *p* < 0.05) of permutation and K-S tests were indicated with square and circle symbols, respectively. **(B)** Total number of transcripts of IL6 family ligands (*IL6, LIF, CLCF1*) expressed in CAFs (*y* axis; error bars denote the 95% confidence interval), stratified by CAF subtype. **(C)** Proportion of malignant subtypes in FOVs with malignant IL6 family receptor expression in the top *versus* bottom quartile of FOVs. **(D)** Spearman correlation between JAK/STAT pathway expression and the expression ratio of *IL6ST* to the sum of *IL6ST* and *IL6R* in malignant cells across all FOVs. Least squares regression line is denoted as a red dotted line, and Spearman rho and p values are annotated. **(E)** Spearman correlation between *GLIS3* expression and the expression ratio of *IL6ST* to the sum of *IL6ST* and *IL6R* for all malignant cells in an FOV, using the subset of FOVs (*n*=155) that were profiled with the augmented panel. Colored dots denote the five FOVs with the greatest absolute number of malignant cells for each subtype, and Spearman rho and p values were annotated. **(F)** The expression ratio of *IL6ST* to the sum of *IL6ST* and *IL6R* in malignant cells (*y* axis; error bars denote the 95% confidence interval), stratified by malignant subtype. \*\*\*\**p* < 0.0001, two-sample Mann-Whitney U test. **(G)** Spatial visualization of *IL6–IL6ST* (*top*) and *IL6–IL6R* (*bottom*) LR interactions between iCAFs and NRP (*left*) or BSL (*right*) cells in representative FOVs. Dots represent ligand gene expression in iCAFs (purple) or receptor gene expression in NRP (green) or BSL (orange) cells. Dot size indicates normalized expression levels and the dot location indicates spatial coordinates of cell centroids. Cells expressing ligands or receptors but not participating in iCAF–NRP (*left*) or iCAF-BSL (*right*) interactions are shown as semi-transparent dots.

Prior studies have identified an association between IL6-producing iCAFs^24^ and EMT,^13,39,40^ which is partially captured by the basal-like subtype.^23,52^ We examined our previously published whole-transcriptome digital spatial profiling (DSP) data on an overlapping cohort of primary tumors, wherein we were able to leverage the whole-transcriptome coverage to sub-classify the ‘basal-like’ state into distinct basaloid, squamoid, and mesenchymal (MES) programs at the tradeoff of multicellular spatial resolution.^6^ This DSP analysis revealed that MES (Spearman *r* = 0.615, *p* < 10^−16^) and NRP (Spearman *r* = 0.283, *p* = 7.66×10^−5^) malignant cells were more spatially co-localized with iCAFs than other malignant subtypes (**Fig. S7C**). Concordant with prior literature,^24,53^ iCAFs demonstrate higher expression of IL6 family ligands compared to myCAFs (*log_2_*FC = 0.442; *p* < 10^−16^, two-sided Mann-Whitney U) (**Fig. 7B; Fig. S7D**), while malignant cells with high expression of IL6 family receptors were enriched for the NRP program (*log_2_*FC = 1.31; *p* = 7.71×10^−5^, two-sided Mann-Whitney U) (**Fig. 7C**). Combined with the SCOTIA results that show a post-treatment enrichment in IL6 family signaling between CAFs and cancer cells (**Fig. 5A-C**), these findings suggest that iCAFs may interact with NRP and MES/BSL cancer cells through IL6 family ligand–receptor signaling pathways in a manner that is enhanced in response to cytotoxic therapy.

Both subunits of the IL6 receptor complex, IL6Rα (*IL6R*) and the signal transducing subunit gp130 (*IL6ST*), can occur in transmembrane and soluble forms. Classical signaling via IL6 binding to membrane-bound IL6Rα and trans-signaling via IL6 forming a complex with soluble IL6Rα and binding to membrane-bound gp130 can have distinct biological effects; the ratio of *IL6ST* to *IL6R* expression determines whether classical or trans-signaling is dominant.^54^ IL6 trans-signaling has faster kinetics and stronger transduction of downstream JAK/STAT pathways than classical signaling,^54^ which is supported in our data by a significant positive correlation between JAK/STAT pathway expression versus the expression ratio of *IL6ST* to the sum of *IL6ST* and *IL6R* (Spearman *r* = 0.409, *p* = 3.96×10^−14^) (**Fig. 7D; Fig. S7E**). Compared to classical signaling, IL6 trans-signaling also leads to greater induced expression of the transcription factor *GLIS3*, which is thought to be a key regulator of the NRP malignant state.^55,56^ In concordance, we observed a positive correlation between *GLIS3* expression, which we included in the augmented panel, and the ratio of *IL6ST* to the sum of *IL6ST* and *IL6R* expression in our SMI data (Spearman *r* = 0.231, *p* = 3.76×10^−3^) (**Fig. 7E**). Furthermore, *IL6ST* was most highly expressed in NRP cells, while *IL6R* was predominantly expressed by BSL cells (**Fig. 7F-G; Fig. S7F**).

## DISCUSSION

In this study, we profiled a cohort of 13 independent neoadjuvant-treated and treatment-naive human pancreatic tumors using high-plex single-cell spatial transcriptomics with a subset featuring matched single-nucleus RNA-sequencing (snRNA-seq) and digital spatial profiling (DSP) data. Combining this unique resource of matched SMI, snRNA-seq and DSP data with a complementary co-culture tumoroid model enabled us to elucidate treatment-associated remodeling of tissue organization and cell–cell interactions in pancreatic cancer. Specifically, we uncovered spatial associations among distinct subtypes of malignant, CAF, and immune cells, revealing differences in cellular architecture between the treated and untreated contexts (**Fig. 2H, 3; Fig. S3**). We observed a spectrum of intraglandular heterogeneity and interspersion with respect to malignant subtypes across patient tumors and individual glands. The predominance of homogeneous glands is consistent with the notion that individual glands are clonal with relatively stable transcriptional states (**Fig. 3; Fig. S3**).^57,58^ However, there was also a substantial number of glands with high intraglandular heterogeneity and interspersion, suggesting that epigenetic differences may exist within the same gland and even among adjacent malignant cells (**Fig. 3; Fig. S3**). Cell type composition analysis uncovered lower abundance of neutrophils in treated specimens (**Fig. 2H**). Lower levels of *CXCL1-3,5,6,8* expression in treated CAFs and malignant cells (**Fig. S2E**) without an associated difference in peripheral neutrophil abundance suggest that chemoradiation therapy may reduce recruitment of neutrophilic immune cells to the tumor.^59^

Existing methods utilizing optimal transport theory to evaluate cell–cell interactions consider potential cell pairs even if they are spatially far apart^60,61^ or depend on a user-defined distance constraint to capture the effect range of ligand diffusivity.^62^ To overcome this limitation, we developed Spatially Constrained Optimal Transport Interaction Analysis (SCOTIA), which extracts spatially adjacent source and target cell clusters, thereby excluding cells that are less likely to communicate. Using SCOTIA, we assessed the strength of interactions across cell type pairs in treated versus treatment-naive tumors and discovered that CAF–malignant interactions were the most enriched in the treated group (**Fig. 4; Fig. S4**). As the most abundant stromal cells in the PDAC TME, CAFs have been shown to promote resistance to chemotherapy and radiotherapy and therefore represent promising therapeutic targets.^31,32,63^ However, treatment-associated signaling between CAFs and malignant cells are not well understood.

Differential expression analysis revealed that more LR pairs were significantly enriched in untreated specimens (*n* = 54) compared to treated specimens (*n* = 22), suggesting there is a baseline level of CAF–malignant cell crosstalk that is disrupted with cytotoxic therapy (**Fig. 5; Fig. S5**). This finding was supported by several lines of evidence as we observed correlated expression of targets downstream of candidate receptors (**Fig. 5F**), and similar expression changes in both human snRNA-seq (**Fig. 5E**) and murine tumoroid co-cultures (**Fig. 6**). A subset of these candidate interactions likely contributes to therapeutic resistance; the underlying mechanisms and translational potential of disrupting such pro-tumorigenic signaling warrants further investigation.

A range of biological pathways were found to be diminished in the treated specimens, including ECM-integrin crosstalk, WNT signaling, interferon signaling (type I and II), and specific cytokine signaling pathways (**Fig. S5C**). Since collagen signaling regulates a diverse set of cellular processes, including cell adhesion, migration, survival and proliferation,^64^ our results suggest that these processes are likely important for the development and progression of pancreatic tumors, and that their disruption is a consequence of cytotoxic treatment. In treated samples, CAF expression of WNT ligands and WNT signaling from CAFs to malignant cells was significantly lower while malignant cell expression of WNT ligands was significantly higher (**Fig. 5D**), consistent with cancer cell acquisition of non-genetic WNT niche independence previously associated with PDAC progression.^65^ These results suggest that autocrine WNT signaling in malignant cells may facilitate escape from cytotoxic therapy-induced cell death.^66,67^

Among the LR pairs enriched in the treated group, several ligands (e.g., FN1, CXCL12, and TGFB1) and receptors (e.g., CXCR3, CXCR4, ACKR3) have been linked to chemoresistance and cancer progression.^33–35,37,68,69^ Several predicted LR pairs that involve TNF, chemokine, and integrin family receptors are non-canonical and not yet fully elucidated. Interestingly, several CAF ligands such as VTN, FN1, and CSF2 were enriched in both untreated and treated specimens but were paired with distinct receptors in these disparate contexts, suggesting that ligands may switch signaling partners because of treatment (**Fig. 5A-B**). In the untreated samples, the receptors paired with these shared ligands were integrins, suggesting a role in cell-matrix and cell-cell adhesion. By contrast, the paired receptors in treated specimens were the TNF family receptor *TNFRSF11B* (osteoprotegerin) and cytokine receptor *CSF3R*, which potentially inhibit TRAIL-activated apoptosis and activate downstream JAK/STAT, NF-κB, and WNT/β-catenin signaling linked to EMT, increased cancer cell survival and invasion, and metastasis.^70,71^

In agreement with prior studies, we observed an increase in iCAF abundance at the expense of myCAFs in treated samples (**Fig. 2G; Fig. S7B**).^6,12^ Inflammatory CAFs express higher levels of IL6 family cytokines (**Fig. 7B; Fig. S7D**), which are associated with EMT, immune evasion, chemo-resistance, and reduced survival.^24,38^ While myCAFs were in close proximity to cancer cells overall (**Fig. 3G**), iCAFs co-localized with NRP and BSL/MES cancer cells specifically and NRP cancer cells also co-localized with CD8 T cells (**Fig. 3F, 7A; Fig. S7A, C**). These findings were consistent with our prior identification of a treatment-enriched spatial neighborhood in PDAC including NRP and MES cancer cells, iCAFs, and CD8 T cells,^6^ which was discovered using whole-transcriptome DSP. It is important to note that in this study, we preferentially selected samples with poor response to neoadjuvant treatment to provide adequate amounts of residual tumor to profile using SMI but this may also contribute to the lack of NRP enrichment in treated samples (**Fig. 2G)**.^6^

The most consistent treatment-enriched candidate LR interaction between CAFs and cancer cells across the SMI, snRNA-seq, and tumoroid co-culture datasets was *CLCF1–CNTFR*, a member of the IL6 cytokine family (**Fig. 5-6**). Moreover, the overall enrichment of chemokine signaling between CAFs and malignant cells (**Fig. 5; Fig. S5**) suggests strengthened intercellular tropism in the post-treatment context. While CAF-derived IL6 and downstream JAK/STAT and MAPK activation has been linked to induction of a mesenchymal cancer cell phenotype in several malignancies including pancreatic cancer^13,39–41^ as well as neuroendocrine differentiation in prostate cancer,^72^ an association with the neural-like progenitor (NRP) malignant cell state has not been previously established. Our SMI data revealed that malignant cells with high expression of IL6 family receptors were enriched for the NRP state (**Fig. 7C**). Moreover, IL6 signaling occurs in two forms: classical signaling describes IL6 binding to the membrane-bound receptor complex consisting of IL6Rα (*IL6R*) and co-receptor gp130 (*IL6ST*), whereas trans-signaling entails IL6 forming a complex with soluble IL6R, which then binds to membranous gp130 to induce stronger signaling through the JAK/STAT and MAPK pathways.^73^ We found that JAK/STAT pathway activation was positively correlated with the expression ratio of *IL6ST* to the sum of *IL6ST* and *IL6R* (**Fig. 7D**), which is consistent with prior studies demonstrating that the ratio of gp130 (*IL6ST*) to IL6Rα (*IL6R*) expression governs the extent of trans-versus classical signaling.^54^ Notably, we discovered a significant monotonic increase in the expression ratio of *IL6ST* to the sum of *IL6ST* and *IL6R* from CLS to BSL to NRP malignant cells. Indeed, IL6 classical and trans-signaling induced a 20% and 65% increase, respectively, in expression of *GLIS3*, a key transcription factor underlying the NRP cell state,^56^ in human airway smooth muscle cells.^55^ Taken together, these results suggest that IL6 family signaling is an important mechanism of interaction between iCAFs and treatment-resistant cancer cells. Heterogeneous expression of *IL6R* and *IL6ST* on cancer cells may drive the balance of classical and trans-signaling to modulate the strength of JAK/STAT and MAPK pathway activation and cancer cell transition towards the BSL/MES or NRP states, which would have important therapeutic ramifications.

This study has several limitations. First, the number of specimens in both the treated and untreated cohorts are relatively small, which somewhat limits the power of the subsequent analyses. Second, lack of pre- and post-treatment matching precluded us from directly inferring the effects of treatment. Third, while the spatial molecular imaging target panel size was large compared to similar contemporary methods, the lack of whole transcriptome coverage restricted the fidelity of cell typing and cell state identification. Fourth, pairwise ligand–receptor interactions likely capture much of but not all the influences a multicellular neighborhood has on an individual cell state. A relatively large number of treatment-associated ligand–receptor pairs were non-canonical, which warrants additional exploration. Future work on cell–cell interactions should extend upon the SCOTIA method to include multicellular influences on an individual cell of interest and model non-ligand–receptor interactions such as those of a mechanical, metabolic, electrical, or junctional nature.

In this study, we integrated novel experimental and computational approaches to enable high-resolution, spatially-guided discovery of treatment-associated remodeling in the pancreatic cancer microenvironment. Further studies are warranted to dissect the specific roles of candidate cell–cell interactions in promoting disease progression and resistance to cytotoxic therapy, and ultimately guide novel therapeutic development. This work provides a broad translational paradigm that leverages single-cell spatial transcriptomics to better understand baseline multicellular neighborhoods/interactions and remodeling of these dynamic relationships under perturbative selection pressure.

## METHODS

### Ethics

All patients in this study were consented without compensation to excess tissue biobank protocol 2003P001289 (principal investigator: C.F.C.; co-investigator: W.L.H.) and used in this research under secondary research protocol 2022P001264 (principal investigator: W.L.H.), which were reviewed and approved by the Massachusetts General Hospital Institutional Review Board. This study was compliant with all relevant ethical regulations.

### Human tumor specimens

Patients selected for inclusion in this study had nonmetastatic pancreatic ductal adenocarcinoma (PDAC) and underwent surgical resection of the primary tumor with or without prior neoadjuvant chemotherapy and radiotherapy (CRT). Most treated patients received several cycles of FOLFIRINOX chemotherapy followed by multi-fraction conformal radiotherapy with concurrent capecitabine or fluorouracil. One patient received additional treatment with losartan (angiotensin II receptor type 1 antagonist; CRTL). Radiotherapy regimens included 30 Gy in 10 fractions, 50.4 Gy in 28 fractions, and stereotactic body radiotherapy 36 Gy in 6 fractions. Serially sectioned formalin-fixed, paraffin-embedded (FFPE) sections (5 µm) of patient-derived primary PDAC tumors were prepared under RNase-free conditions. Sections were attached to the non-charged side of Superfrost Plus Micro Slides (VWR).

### Custom probe selection

We designed 30 custom probes to target malignant subtypes described in Hwang et al.^6^ For each malignant subtype, we developed a backward stepwise regression model (leaps v3.1, R v4.1.0) to determine gene sets with the highest predictive value and excluded genes with higher Pearson correlation with off-target malignant subtypes. We leveraged a combination of the regression model results, well-known transcription factors and canonical marker genes to select the 30 target genes.^6,23,52,56^

### Sample preparation for spatial molecular imaging

To prepare the tissue samples for spatial molecular imaging on a pre-commercial SMI platform (NanoString Technologies Inc, Seattle, WA), slides with PDAC sections were baked overnight at 60°C to ensure FFPE tissue adherence to the glass slides. Samples underwent deparaffinization, proteinase K (3 μg ml^−1^; ThermoFisher) digestion, and heat-induced epitope retrieval (HEIR) procedures to expose target RNAs and epitopes using the Leica Bond RX system. Proteinase K incubation at 40°C for 30 min and HEIR at 100°C for 15 min in Leica buffer ER1 conditions were used. The PDAC samples were rinsed with diethyl pyrocarbonate (DEPC)-treated water (DEPC H_2_O) twice before incubating with 1:1000 diluted fiducials (Bangs Laboratory) in 2X SSCT (2X saline sodium citrate, 0.001% Tween 20) solution for 5 min at room temperature. Excess fiducials were removed by rinsing the samples with 1X phosphate buffered saline (PBS), followed by fixation with 10% neutral buffered formalin (NBF) for 5 min at room temperature. Fixed samples were rinsed with Tris-glycine buffer (0.1M glycine, 0.1M Tris-base in DEPC H_2_O) and 1X PBS for 5 min each before being blocked using 100 mM N-succinimidyl acetate (NHS-acetate, ThermoFisher) in NHS-acetate buffer (0.1M NaP, 0.1% Tween pH 8 in DEPC H_2_O) for 15 min at room temperature. Prepared samples were rinsed with 2X saline sodium citrate (SSC) for 5 min and then a Adhesive SecureSeal Hybridization Chamber (Grace Bio-Labs) was placed to cover the samples.

RNA ISH probes (960-plex for the base panel, 30 additional probes for the custom panel, and 19 negative control probes) were denatured at 95°C for 2 min and then placed on ice before preparing the ISH probe mix (1 nM ISH probes, 1X Buffer R, 0.1 U μl^−1^ SUPERaseIn™ in DEPC H_2_O). The ISH probe mix was pipetted into the hybridization chamber and the chamber was sealed using adhesive tape. Hybridization was performed at 37°C overnight to prevent evaporation. After the overnight hybridization, samples were washed twice with 50% formamide (VWR) in 2X SSC at 37°C for 25 min, rinsed twice with 2X SSC for 2 min at room temperature, and then blocked with 100 mM NHS-acetate for 15 min. After blocking, the samples were washed twice using 2X SSC for 2 min at room temperature. Custom-made slide covers were attached to the sample slide to form a flow cell.

### Data acquisition for spatial molecular imaging

Target RNA readout on the SMI instrument was performed, following published protocols.^19^ In brief, the assembled flow cell was loaded onto the SMI instrument and the tumor samples were washed with a reporter wash buffer to rinse the samples and remove air bubbles. The reporter wash buffer consisted of 1X SSPE, 0.5% Tween 20, 0.1 U μl^−1^ SUPERase•In RNase Inhibitor (20 U μl^−1^), 0.1% Proclin 950 and DEPC H_2_O. Once the flow cell was loaded onto the instrument, the entire flow cell was scanned and 15-25 fields of view (FOVs) were placed on each slide for RNA readout. The first RNA readout cycle was initiated by flowing 100 µl of reporter pool 1 into the flow cell and incubating for 15 min. After incubation, 1 ml of reporter wash buffer was passed across the flow cell to wash out the unbound reporter probes, followed by replacing reporter wash buffer with imaging buffer (80 mM glucose, 0.6 U ml^−1^ pyranose oxidase from *Coriolus* sp., 18 U ml^−1^ catalase from bovine liver, 1:1,000 Proclin 950, 500 mM Tris-HCl buffer pH 7.5, 150 mM sodium chloride and 0.1% Tween 20 in DEPC H_2_O) prior to imaging. Eight to nine Z-stack images (0.8 um step size) of each FOV were acquired and then fluorophores on the reporter probes were UV cleaved and washed off with strip wash buffer (0.0033× SSPE, 0.5% Tween 20 and 1:1000 Proclin 950). This fluidic and imaging procedure was then repeated for the remaining 15 reporter pools, and the 16 cycles of reporter hybridization-imaging were repeated 8 times to increase RNA detection sensitivity.

After 9 complete rounds of RNA readout, PDAC samples were incubated with a fluorophore-conjugated antibody cocktail against CD298 (Abcam, EP1845Y), B2M (Abcam, EP2978Y), PanCK (Novus, AE-1/AE-3), CD45 (Novus, 2B11 + PD7/26), and CD3 (Origene, UMAB54) proteins and DAPI in the same instrument for 1 hr. Eight to nine Z-stack images for 5 channels (5 antibodies and DAPI) were captured after unbound antibodies and DAPI were washed with reporter washing buffer and the flow cell was filled with imaging buffer.

### Image processing and cell segmentation

Raw image processing and feature extraction were performed using an in-house SMI data processing pipeline^19^ which includes registration, feature detection, and localization. 3D rigid image registration was made using fiducials embedded in the samples matched with the fixed image reference established at the beginning of the SMI run to correct for any shifts. Secondly, the RNA image analysis algorithm was used to identify reporter signature locations in the *x*, *y*, and *z* axes along with the associated confidence. The reporter signature locations and features were collated into a single list. Lastly, the *xyz* location information of individual target transcripts was extracted and recorded in a table by a secondary analysis algorithm, as described in He et al.^19^

The Z-stack images of nuclear (DAPI) and CD298/B2M (surface) staining were used for drawing cell boundaries on the samples. A cell segmentation pipeline using a machine learning algorithm^74,75^ was used to accurately assign transcripts to cell locations and subcellular compartments. The transcript profile of individual cells was generated by combining target transcript location and cell segmentation boundaries. Cells with fewer than 60 total transcripts were omitted from downstream analysis. Expression profiles were normalized for each cell by dividing its raw count vector by total counts and multiplied by the scale factor 10,000 before log transformation.

### Cell type annotations

Cell type identification was performed with a supervised clustering method, Insitutype.^21^ High-confidence malignant and non-malignant cells were first selected as anchors based on protein (PanCK) and expression (*KRT5*, *KRT7*, *KRT8*, *KRT18*, *KRT19*, and *KRT20*) of cytokeratin marker genes. Cells with high levels of cytokeratin protein and RNA expression were defined as malignant anchors, whereas cells exhibiting low expression levels were selected as non-malignant anchors. Anchor cells were further filtered by removing outliers not near cluster centroids based on K-means clustering. The averaged gene profiles of anchor cells were used as references for Insitutype to classify all cells into malignant and non-malignant categories. Batch correction was performed on all non-malignant cells by treating each tissue slide as a batch using ComBat.^76^ Subsequently, corrected non-malignant anchor cells were classified into cell types using reference profiles derived from a whole transcriptome snRNA-seq dataset consisting of 43 primary human PDAC specimens.^6^ To obtain reference profiles for each cell type, we calculated the average gene expression of cells classified as a given type in the snRNA-seq dataset. A SMI-derived reference profile was established on those anchor cells by averaging gene expression within each cell type that was subsequently used for assigning all non-malignant cells into known cell types. Clustered cells were displayed using the runUMAP function in Giotto.^77^

To further subtype CAFs and malignant cells, representative genes for some subtypes^6,24,25^ were selected for supervised sub-clustering (**Fig. 2B**). To leverage the 30 custom probes that were designed for annotating malignant subtypes, we first performed supervised clustering on the subset of samples (*n* = 6) that were profiled with these additional custom probes (990-plex panel) to generate modified reference profiles for the subtypes of interest, which we then applied to cluster the remaining samples (*n* = 9; 960-plex panel), including two samples that were analyzed by both panels.

### Malignant gland extraction and classification

To extract malignant glands, we aggregated two independent clustering assignments that were determined using Density-Based Spatial Clustering of Applications with Noise (DBSCAN)^29,30^ algorithm (from the *sklearn* package in Python) on the pan-cytokeratin (PanCK) protein antibody channel and on the spatial locations of malignant cells.

When performing DBSCAN on the PanCK protein antibody channel (whose pixels were downsampled by a factor of 19 for reduced computing time), we set the maximum distance between neighboring PanCK-positive pixels to 60 pixels (10.8 µm) and the minimum number of PanCK-positive pixels in a neighborhood to 2. We leveraged the immunofluorescence (IF)-based clustering results to determine cluster assignments for each malignant cell by identifying the nearest PanCK-positive pixel. If the nearest PanCK-positive pixel was within 180 µm, then we labeled that cell with the same cluster as the positive pixel. Otherwise, the malignant cell was not assigned to any malignant gland cluster. Four out of 257,767 malignant cells (0.00155%) were not assigned to a gland using the IF-based approach.

When performing DBSCAN on the locations of malignant cells as defined by spatial transcriptomics, we set the maximum distance between neighboring cells to 20 pixels (3.6 µm) and the minimum number of cells in a neighborhood to 2. Some malignant cells were not assigned to any glandular cluster. Seven out of 257,767 malignant cells (0.00272%) were not assigned to a gland using the spatial location-based approach.

We then acquired a consensus between the two clustering assignments. If two distinct clusters in one clustering method have at least one cell in the same cluster by the other method, then these two clusters are merged. The parameters in the two individual approaches were chosen to over-cluster the malignant cells, which was then compensated for using this consensus method. Extracted malignant glands with less than 5 cells were disbanded, yielding a total of 9,017 malignant cells (3.50%) that were not assigned to a gland.

Only extracted malignant glands with less than 500 cells were included in downstream glandular analyses. We classified a malignant gland as predominantly consisting of a certain malignant subtype if one of the subtypes comprised at least 70% of the cells in the gland. To identify malignant gland size differences among subtypes, we shuffled the subtype annotations across all malignant cells within each FOV to generate a null distribution. We performed 1,000 permutation tests to obtain the significance. When comparing average gland sizes between malignant subtypes, we reported significance values determined by the two-sided Kolmogorov-Smirnov test.

### Glandular subtype heterogeneity and interspersion

The subtype heterogeneity *H_g_* of a malignant gland *g* was computed as:

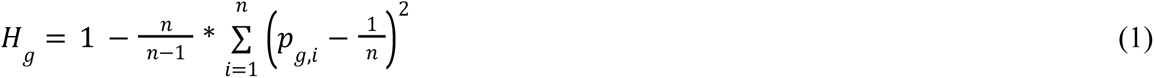

where *p_g,i_* is the proportion of each subtype *i* in gland *g* and *n* is the number of subtypes (*n* = 3 in this study). We measured the interspersion *I_g_* of subtypes within a malignant gland *g* to quantify the difference between glands composed of a mostly alternating pattern of subtypes and glands composed of larger clusters of the same subtype:

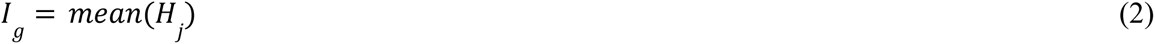

For each malignant cell in the gland, we first computed the subtype heterogeneity *H_j_* of the cell of interest and its neighborhood *j*, which we defined to be any malignant cell within 10 μm in the same gland. We then determined the interspersion *I_g_* for each malignant gland *g* by computing the mean subtype heterogeneity across cells that have at least one neighbor.

We performed 1,000 permutation tests to determine whether the observed heterogeneity and interspersion values differed significantly from the null distribution. The null distribution was established by shuffling malignant subtype annotations within each FOV and recomputing the heterogeneity and interspersion of each gland. When comparing the observed and expected distributions, we reported significance values determined by the two-sided Kolmogorov-Smirnov test.

Glands were classified as having high heterogeneity and/or interspersion if their heterogeneity and/or interspersion scores were in the top quartile of glands across all non-permuted samples. Similarly, glands were classified as having low heterogeneity and/or interspersion if their scores were in the bottom quartile.

### Exponential model

We developed an exponential-based model to compare the pairwise proximity of different cell types and subtypes. The exponential decay captured by this model follows the assumption that the interaction between two cells rapidly declines as their pairwise distance increases. To compute the proximity between cell types *A* and *B*, we first measured the distances between each cell type pair within each FOV, and we then transformed the measured distances using the following exponential function:

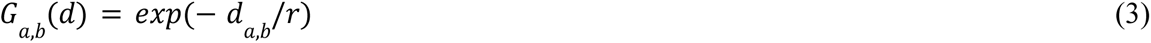

where *d*_a,b_ is the distance between cells *a* and *b* (of types *A* and *B*, respectively) and *r* is the decay radius in μm. For most analyses, we presented results using a decay radius of *r =* 50 μm, but our findings were robust across varying *r*. For each cell *a*, we summed the exponential-transformed distance measurements across all cells *b* in the same FOV. This yielded an observed distribution of summed transformed values that we compared to the expected null distribution.

We performed 1,000 permutations to generate a null distribution, which was used to determine whether the observed mean summed transformed values differed significantly from their expectation. A permutation test was necessary to control for differences in pairwise proximities that are a result of variations in cell density across malignant subtypes. The null distribution was acquired by shuffling the cell type annotations within major cell type categories (e.g. malignant, CAF, and immune) within each FOV. This allowed us to maintain the high level attributes of the tumor architecture in this analysis. For example, when comparing the proximities between malignant subtypes and CD8 T cells, we generated an expected distribution by shuffling the malignant subtype annotations. Similarly, when comparing the proximities between CAF subtypes and malignant cells, we generate an expected distribution by shuffling the CAF subtype annotations. In addition to permutation tests, we also used the Kolmogorov-Smirnov (K-S) test to measure the difference between the expected distribution *D*_exp_ and the observed distribution *D*_obs_ of summed transformed values.

We computed a statistic *t* such that *P*(*D_exp_* < *t*) = *q*. For the majority of analyses, we presented results using a quantile threshold of *q =* 0.95, but our findings were robust across varying *q*. We then measured the proportion of summed transformed values that were greater than *t*. This proportion was used to compute the log_2_ fold change between the expected and observed distributions for each permutation. We performed two-sided Mann Whitney U tests to compare the log_2_ fold changes between treated and untreated cohorts.

Of note, the quantity obtained from the exponential-based model is asymmetric. Above, we describe a cell type *A*-centric model, in which we acquired an observed distribution by measuring the pairwise distance between each cell *a* and all other cells *b* in the same FOV. In a cell type *B*-centric model, we can obtain a distinct observed distribution by summing the transformed distance measures between each cell *b* and all other cells *a* in the same FOV.

### Cell density classifier

Cell density was calculated as the total number of cells of each subtype, plus a pseudocount of one, divided by the area of each FOV. A *k* nearest neighbor classifier using the densities of NRP, CLS, BSL, iCAF, myCAF, neutrophil, natural killer, macrophage, and endothelial cells was trained while withholding all of the FOVs from one of the samples. The density vectors were concatenated, log-transformed, and then the first *P* principal components were used as features. The reported accuracy is the fraction of correctly assigned FOVs in each withheld sample, iterated and averaged across all sample combinations. To ensure robustness, we employed a consensus strategy, wherein we considered *P* = 6 to 12 as well as *k* = 21 to 51 neighbors.

### Topological data analysis and K-nearest neighbor classifier

For each cell type, we first normalized spatial cell positions such that *x*- and *y*-coordinates were in the range [0, 1]. Second, we calculated the persistence diagram based on Euclidean distances. Finally, the persistence images^28^ were calculated using a one-dimensional image with 5 pixels and a standard deviation of 0.01. The pixel vectors were concatenated, log-transformed, and then the first *P* principal components were used as features to train a *k* nearest neighbor classifier. Training and testing splits were performed in the same way as for the cell density classifier. To ensure robustness, we employed a consensus strategy and we considered *P* = 6 to 12 as well as *k* = 21 to 51 neighbors.

### Spatially Constrained Optimal Transport Interaction Analysis (SCOTIA)

Ligand–receptor (LR) interactions were modeled using an unbalanced optimal transport (OT) algorithm.^60,78^ We included LR pairs from the FANTOM5 database,^79^ which has been widely used to predict cell–cell interactions in scRNA-seq analysis,^51^ and only secreted ligands were included based on the Human Protein Atlas.^80^ In total, 422 LR pairs were covered in our SMI panels. To infer LR interaction strength between cell type pairs, we spatially constrained the potential search space of communicating pairs by selecting spatially adjacent source and target cell clusters using DBSCAN.^29,30^ Since cell densities varied across FOVs and samples, the hyperparameter *eps* for maximum distance between neighboring cells within each cluster was dynamically estimated for each cell type in each FOV by finding the maximum overlap of cluster labels between DBSCAN (using Python package *sklearn*) and the Forest Fire clustering algorithm.^81^ Briefly, the hyperparameter *eps* in the DBSCAN algorithm was tuned in the range of 15 to 150 with a step size of 5, while the hyperparameter temperature *c* in the Forest Fire clustering algorithm was adjusted in the range of 1 to 60 with a step size of 1. The adjusted Rand index scores^82^ between the two clustering methods with different hyperparameter values were calculated using the *adjusted_rand_score* function from the Python package *sklearn*. The *eps* that had the highest adjusted rand index score was selected as the most optimal *eps*. The minimum number of cells per cluster was set at 10. Clusters were considered ‘adjacent’ when the shortest distance between two cells in each cluster was less than 50 µm.

We then inferred LR interaction scores for cell pairs between spatially adjacent clusters by minimizing the total cost for transporting ligand gene expression to receptor gene expression, which was formulated as:

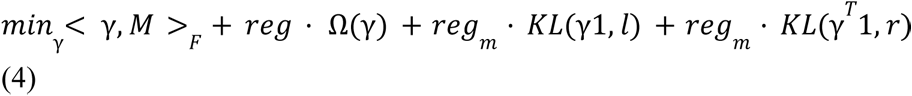

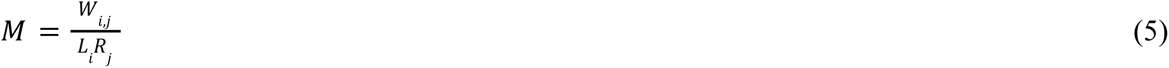

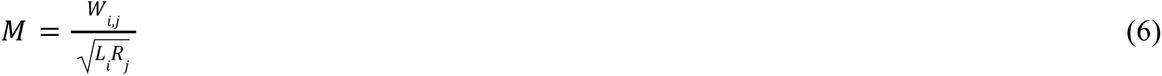

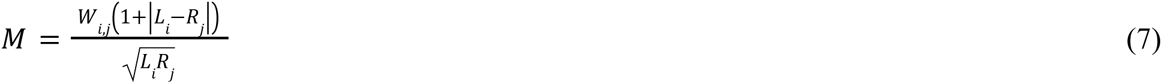

Where <γ, ***M***>*_F_* is the Frobenius inner product of γ and ***M***. γ (≥ 0) is the transport plan and is considered the interaction score; ***M*** is one of the three customized cost matrices (equations 5-7), which integrate spatial distance with expression distance: *W_i,j_* is the spatial distance between source cell *i* and target cell *j*; *L_i_*and *R_j_* are the normalized expression of the ligand gene in cell *i* and the receptor gene in cell *j*. Ω is an entropy term penalizing one-to-one transport cases; *reg* is the entropy regularization term, the default is 1; *reg*_m_ is the marginal relaxation term, the default is 2; ***l*** and ***r*** are the distribution of ligand gene expression in source cells and the receptor gene expression in target cells, respectively; **1** is a vector of ones; *KL* is the Kullback-Leibler divergence and the two *KL* terms are used to relax the difference in total expression between ligand and receptor genes.

After computing LR interaction scores, we performed permutation tests to identify robust LR pairs between any given cell type pair. To maintain the original cell type composition in spatial neighborhoods, the null distribution was established by shuffling gene expression and randomizing cell locations within a small range (from −20 to 20 µm). The number of permutations was 50. To identify differentially enriched LR interactions between the treated and untreated conditions, we performed two-sided Mann-Whitney U tests and constructed a mixed effects model with the sample ID as a random effect and treatment status as the fixed effect covariate. The mixed effects model was implemented using the *mixedlm* function from the Python package *statsmodels*. Multiple testing correction was performed using the Benjamini-Hochberg approach.^83^

### Ligand–receptor target gene and gene set enrichment analysis

The correlation between inferred LR interactions and the expression of downstream target genes was investigated by integrative analyses with the KEGG^48^ and TRRUST v2^49^ databases. Ligand–receptor–transcription factor (TF) pathways were extracted from the KEGG database with the Python package *bioservices*.^84^ Activated or repressed target genes of the TFs were extracted from the TRRUST v2 database.^49^ To apply this approach to our SMI data, we estimated the gene expression correlation (Spearman) between receptors and their target genes; the average expression of the top 10 (for activated genes) and bottom 10 (for repressed genes) highly-correlated targets was computed for each LR pair. Pathways enriched in association with up- or down-regulated ligands and receptors were obtained by Gene Ontology (GO) enrichment analysis.^85^

### Murine stromal tumoroid co-cultures

To establish 3D murine CAF-cancer cell co-cultures, 3000 dissociated ductal cells (KP; *Kras*^LSL-G12D/WT^;*Trp53*^FL/FL^) that were infected with adenovirus bearing Cre recombinase (Ad5-CMV-Cre) and genotyped at the *Kras* and *Trp53* loci to ensure proper recombination^86^ were combined with 24,000 pancreatic stellate cells (PSCs; PSC4 line^24^) and seeded in 20 μl Matrigel domes in a 24 well plate and covered with 1 ml of DMEM+10% FBS. After 24 hours, 5-fluorouracil (5-FU; final concentration 3 μM) or vehicle (DMSO) was added to all wells. On days 4 and 6, half of the media was replaced with fresh media containing 5-FU or vehicle to achieve the same final concentrations. On day 7, the co-culture tumoroids were digested in TrypLE for 30 minutes at 37°C, then washed with PBS three times. The resulting pellet was subjected to nuclei isolation as described for organoids in Hwang et al.^6^

### Single-nucleus RNA-sequencing

The isolated nuclei of each sample (5-FU vs vehicle) were processed using the fixation protocol provided by Parse Biosciences for their Evercode^TM^ single-cell technology with minor modifications in centrifugation as follows: spins were done at 350 x *g* at 4°C for 10 minutes with acceleration speed 9 and deceleration speed 5. Up to 4 million nuclei were transferred into a 15 mL conical pre-coated with 5% BSA and spun down. The pellet went through a series of washes and filtering, as detailed in the manufacturer protocol, and was ultimately aliquoted and slowly frozen in the −80°C freezer for storage.

The Evercode Whole Transcriptome Mini (Parse Biosciences) protocol was closely followed for barcoding and library preparation. In brief, the nuclei were quickly thawed in a 37°C water bath and counted on a hemocytometer. The loading schematic provided by Parse was modified in that we increased the “Max Number Barcoded Cells” to 25K to account for losses. Single nuclei were barcoded through iterative rounds of ligation and split/pooling using Round 1, Round 2, and Round 3 Plates. The barcoded nuclei were counted on a hemocytometer and 10K cells were loaded into PCR tubes for each sub-library, adding Dilution Buffer as needed to bring the final volume to 25 μl. After amplification of the barcoded cDNA, each sublibrary underwent fragmentation, end repair, A-tailing, adaptor ligation, and index PCR in preparation for sequencing, as detailed in the manufacturer protocol. Notably, we performed 5 cycles for the first round of cycling and 8 cycles for the second round of cycling during cDNA amplification, and 13 cycles for index amplification based on the recommendations listed in the manufacturer protocol. The library was sequenced using a NovaSeq SP flowcell with the following parameters: read type = paired end, fragment size = 400, length of read 1 = 208, length of read 2 = 86, length of index = 6, PhiX spike-in = 5%.

Raw Illumina sequencing reads were converted into FASTQ files using the bcl2fastq conversion software. The FASTQ files were preprocessed using the standard Parse pipeline and the annotated reference genome of GRCm39 (release version 108). Default parameters in the pipeline were used. A matrix containing the raw counts of gene expression across individual nuclei exceeding the minimum transcript threshold (840 transcripts for treated cells; 835 transcripts for untreated cells) was obtained. The minimum transcript threshold was determined as the steepest point in the barcode-rank plot, which orders the transcript counts per nucleus along log transformed axes.

The dataset was transferred into Seurat for further analysis. Low quality cells were removed if the cell expressed: (a) more than 6,000 genes, (b) more than 20,000 transcripts, or (c) more than 15% mitochondrial transcripts. The data was then normalized using the Seurat NormalizeData function with default parameters. The top 50 principal components and the Seurat Integration function (with default parameters) were used to correct for batch effects across treated and untreated groups, cluster cells and compute UMAP reductions. Identified clusters were annotated as malignant (*EPCAM*, *KRT7*, *CDH1*, *KRT18*, *KRT19*) or CAF (*COL1A1*, *COL1A2*, *FGF7*, *FN1*, *POSTN*, *VCAN*) using a subset of canonical epithelial and fibroblast marker genes. Clusters were additionally separated into treated and untreated conditions.

## DATA AND CODE AVAILABILITY

The raw and processed SMI and co-culture snRNA-seq data are available at Mendeley Data (https://doi.org/10.17632/kx6b69n3cb.1) and Zenodo (https://doi.org/10.5281/zenodo.7963531). The snRNA-seq and GeoMx datasets^6^ are available from GEO: GSE202051 and GSE199102, respectively. The SCOTIA package and our analysis code have been uploaded to Zenodo (https://doi.org/10.5281/zenodo.7963527).

## ACKNOWLEDGEMENTS

We are grateful to the patients and families who contributed their time and surgical specimens to this study. We thank K. Cormier for assistance with tissue sectioning; E. Miller, P. Danaher, S. He, S. Wiegel, A. Wardhani, E. Rueckert from NanoString for assistance with project coordination and data analysis; and D. Moschella, T. Balducci, S. McSorley, S. Sullivan, M. Pivovarov, K. Yee, K. Mercer, J. Teixeira, and K. Anderson for administrative and technical support. This work was supported in part by NCI K08CA270417 (W.L.H.), Burroughs Wellcome Fund Career Award for Medical Scientists (W.L.H.), Pancreatic Cancer Action Network Career Development Award (W.L.H.), SU2C-Lustgarten Foundation (T.J., T.S.H., D.T.T.), Robert L. Fine Cancer Research Foundation (D.T.T.), the Evergrande Center (J.C., M.H.), and the Helmsley Foundation (J-W.C., M.H.). The funders had no role in study design, data collection and analysis, decision to publish or preparation of the manuscript.

## Contributions

J.C., C.S., M.H., and W.L.H. developed the study concept. T.K.K., Y.K., N.S., and J.M.B. performed spatial molecular imaging using a panel including custom probes designed by C.S. and W.L.H.. J.C., C.S., M.T.G., J-W.C., P.L.W., J.W.R., M.H., and W.L.H. analyzed and interpreted the spatial molecular imaging data. J.C. and M.H. developed the SCOTIA method with input from C.S. and W.L.H.. S.W., J.S., J.A.G., X.Y., D.G., P.L.W., N.A.L., and W.L.H. designed and performed the *in vitro* experiments and single-nucleus RNA-sequencing. J.L.B. and M.M-K. guided histological sectioning and staining. M.Q., T.S.H., J.Y.W., H.R., C.F.C., and M.M-K. provided clinical insights and access to patient specimens. J.C., C.S., and M.T.G. generated the tables and figures with guidance from M.H. and W.L.H.. Funding for the work was provided by W.L.H., D.T.T., T.J., and M.H. The study was supervised by W.L.H., M.H., D.T.T., T.J., and R.W.. J.C., C.S., P.L.W., M.H., and W.L.H. wrote the manuscript, and all authors reviewed the manuscript.

## SUPPLEMENTARY FIGURES

**Fig. S1. Spatial molecular imaging experimental workflow and probe set. (A)** Schematic of the spatial molecular imaging RNA assay workflow. RNA targets in the FFPE tissue slide that are bound to *in situ* hybridization (ISH) probes are subject to cyclic readout of 16 sets of reporters conjugated to four different fluorophores, which bind to the different reporter-landing domains on the ISH probes. High-resolution images are acquired during each round of reporter hybridization. Fluorophores are then UV cleaved and washed off the reporters before the slide is incubated with the next set of reporters. **(B)** Gene overlap between the seven malignant lineage programs,^6^ the base 960-plex probe set and the 30 custom probes (color legend).

**Fig. S2. Cell type composition across untreated and treated pancreatic cancer samples. (A)** UMAP showing subsets of vascular, lymphoid, and myeloid cells. **(B)** Proportions of major cell types (from **Fig. 2C-D**) across untreated and treated tumors with (*left*) or without (*right*) malignant cells included. U, untreated; T, treated; b, base 960-plex panel; a, augmented 990-plex panel. **(C)** Comparison of malignant/non-malignant cell numbers in treated and untreated FOVs. Pairwise comparisons were performed using the two-sided Mann-Whitney U test (\*\*\**p* < 0.001; ns, not significant). **(D)** Proportions of major non-malignant cell types in treated and untreated FOVs (\*\*\**p* < 0.001; ns, not significant); *p* values were adjusted with Benjamini-Hochberg correction. **(E)** Expression of chemokines with a role in neutrophil recruitment (CXCL1/2/3/5/6/8) in treated/untreated CAFs and malignant cells (\*\**p* < 0.01; \*\*\**p* < 0.001; ns, not significant); *p* values were adjusted with Benjamini-Hochberg correction.

**Fig. S3. Glandular heterogeneity and multicellular neighborhoods in pancreatic cancer. (A)** Distribution of the number of cells across malignant glands. **(B)** Observed (black) and expected (gray) distributions of the interspersion of malignant glands, subset to glands in the top quartile of heterogeneity. Statistical test legend shared with panels D, G and H. **(C)** Three-dimensional depiction of the summed exponential functions (*z* axis height and color bar, decay radius *r* = 50 μm) that are generated from CD8 T cells for a representative FOV, with spatial locations of shuffled malignant subtype annotations shown as colored dots, which serves as the null distribution. **(D)** Log_2_ fold change (*y* axis; error bars denote the 95% confidence interval) between the observed and expected summed exponential-transformed distances between malignant subtypes and CD8 T cells (using a malignant-centric model), for varying quantile thresholds (*x* axis). **(E-F)** Three-dimensional topological depiction of the summed exponential functions that are generated from CD8 T cells for varying decay radii *r* (in μm) **(E)** and that are generated from malignant subtypes, using decay radius *r* = 50 μm **(F)**. **(G-H)** Log_2_ fold change (*y* axis; error bars denote the 95% confidence interval) between the observed and expected summed exponential-transformed distances between malignant subtypes and CD8 T cells (using a malignant-centric model on the *left* and a CD8 T cell-centric model on the *right*) **(G)** and between CAF subtypes and malignant cells (using a CAF-centric model on the *left* and a malignant-centric model on the *right*) **(H)** across varying decay radii (*x* axis).

**Fig. S4. The permutation test strategy used in SCOTIA. (A)** Pearson correlation of receptor gene expression between malignant subtypes or ligand gene expression between CAF subtypes. **(B)** Schematic of the permutation test model used for spatial molecular imaging data. The null distribution was established by randomizing cell locations within a small range (from −20 to 20 µm) (*top*) while shuffling gene expression (*bottom*) in each FOV. **(C)** Neighborhood composition for each cell type from one example FOV with the original (*top*), permuted (*middle*) and shuffled negative control (*bottom*) data. The negative control was constructed by shuffling cell type labels without any constraints. Neighborhood cells were defined as cells within a radius of 30 µm. **(D)** The top five strongest interacting cell type pairs inferred by using cost function equation 6 (*left*) and 7 (*right*) (**Methods**). Dot size represents the number of permutation test-significant LR pairs, colored based on the average LR interaction score. Bar plot indicates the average interaction strengths of each cell type pair for the treated and untreated groups. U, untreated; T, treated; b, base 960-plex panel; a, augmented 990-plex panel.

**Fig. S5. Pathway enrichment and target gene analysis for the SMI data. (A)** Volcano plots showing the log_2_ fold change of ligand–receptor (LR) interaction scores between treated and untreated samples (*x* axis) versus –log_2_ adjusted *p* value (*y* axis) with CAFs as source and malignant cells as target, related to Fig. 5A. The cost functions used were equation 6 (*left*) and 7 (*right*) (**Methods**). **(B)** Volcano plot showing the differentially enriched LR interactions inferred with a mixed effects model (**Methods**). **(C)** Pathway enrichment analysis with ligand (*left*) or receptor (*right*) gene sets from Fig. 5B. Top pathways enriched in untreated (purple) and treated (red) tumors are shown. **(D)** Volcano plots showing the treatment-enriched/depleted LR interactions between five other abundant cell type pairs. Boxplots summarizing the difference in target gene expression between significantly up- and down-regulated LR groups. Two-sided Mann-Whitney U test (\**p* < 0.05; \*\**p* < 0.01; \*\*\**p* < 0.001; ns, not significant).

**Fig. S6. Pathway enrichment analysis for the co-culture tumoroid data. (A)** Pathway enrichment analysis with the ligand gene set (from CAFs, *left*) or receptor gene set (from malignant cells, *right*) that were significantly higher (red) or lower (purple) in the treated versus treatment-naïve co-culture tumoroids, related to Fig. 6C.

**Fig. S7. Association between IL-6 family signaling, inflammatory CAFs and specific malignant subtypes. (A)** Log_2_ fold change (*y* axis; error bars denote the 95% confidence interval) between the observed and expected summed exponential-transformed distances between malignant subtypes and iCAFs (using a malignant-centric model on the *left* and an iCAF-centric model on the *right*) across varying decay radii (*x* axis). **(B)** Proportions of CAF subtypes per FOV (*n* = 320) in SMI (*left*) and per sample (*n* = 43) in single-nucleus RNA-seq^6^ (*right*) datasets, stratified by treatment. **(C)** Proportion of iCAFs and malignant subtypes per FOV in whole-transcriptome digital spatial profiling data (DSP; *n* = 21).^6^ Least squares regression line was denoted as a red dotted line, and Spearman rho and p values were annotated. **(D)** Number of transcripts of specific IL6 family ligands expressed in CAFs (*y* axis; error bars denote the 95% confidence interval) in the SMI dataset, stratified by CAF subtype. **(E)** Spearman correlation between JAK/STAT pathway expression and *IL6R* (*left*) and *IL6ST* (*right*) expression in malignant cells. Least squares regression line was denoted as a red dotted line, and Spearman rho and p values were annotated. **(F)** *IL6R* (*left*) and *IL6ST* (*right*) expression in malignant cells (*y* axis; error bars denote the 95% confidence interval) in the SMI dataset, stratified by malignant subtype. \*\*\*\**p* < 0.0001, two-sample Mann-Whitney U test.

## SUPPLEMENTARY TABLES

**Table S1**

Clinicopathological parameters and quality control metrics for each sample.

**Table S2**

Overlapping and unique ligand-receptor pairs inferred by using different databases.

**Table S3**

Ligand receptor pairs used for SCOTIA analysis.

