## Supplementary figures and images for "Therapy-associated remodeling of pancreatic cancer revealed by single-cell spatial transcriptomics and optimal transport analysis"

### Supplementary Figures

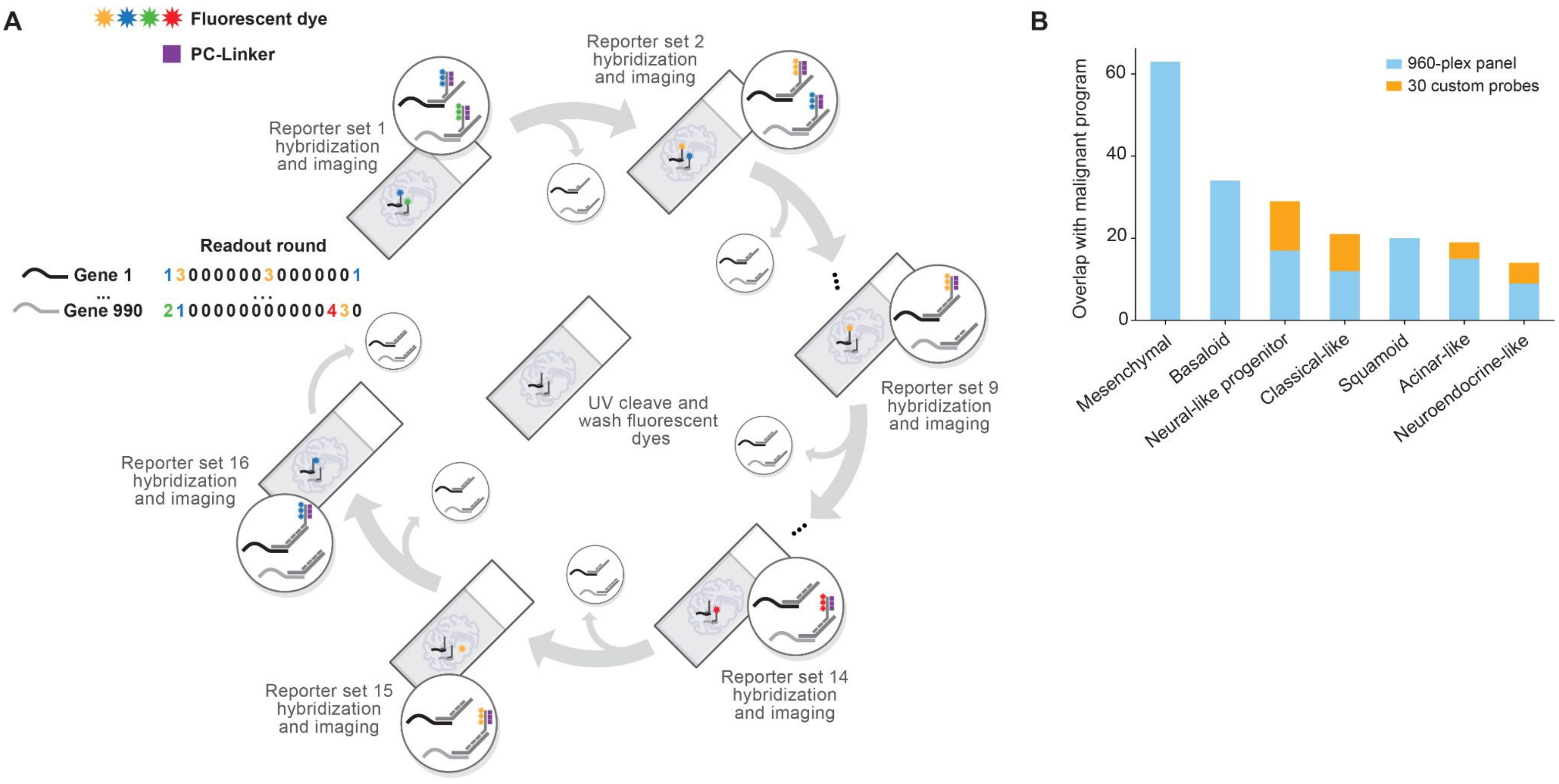

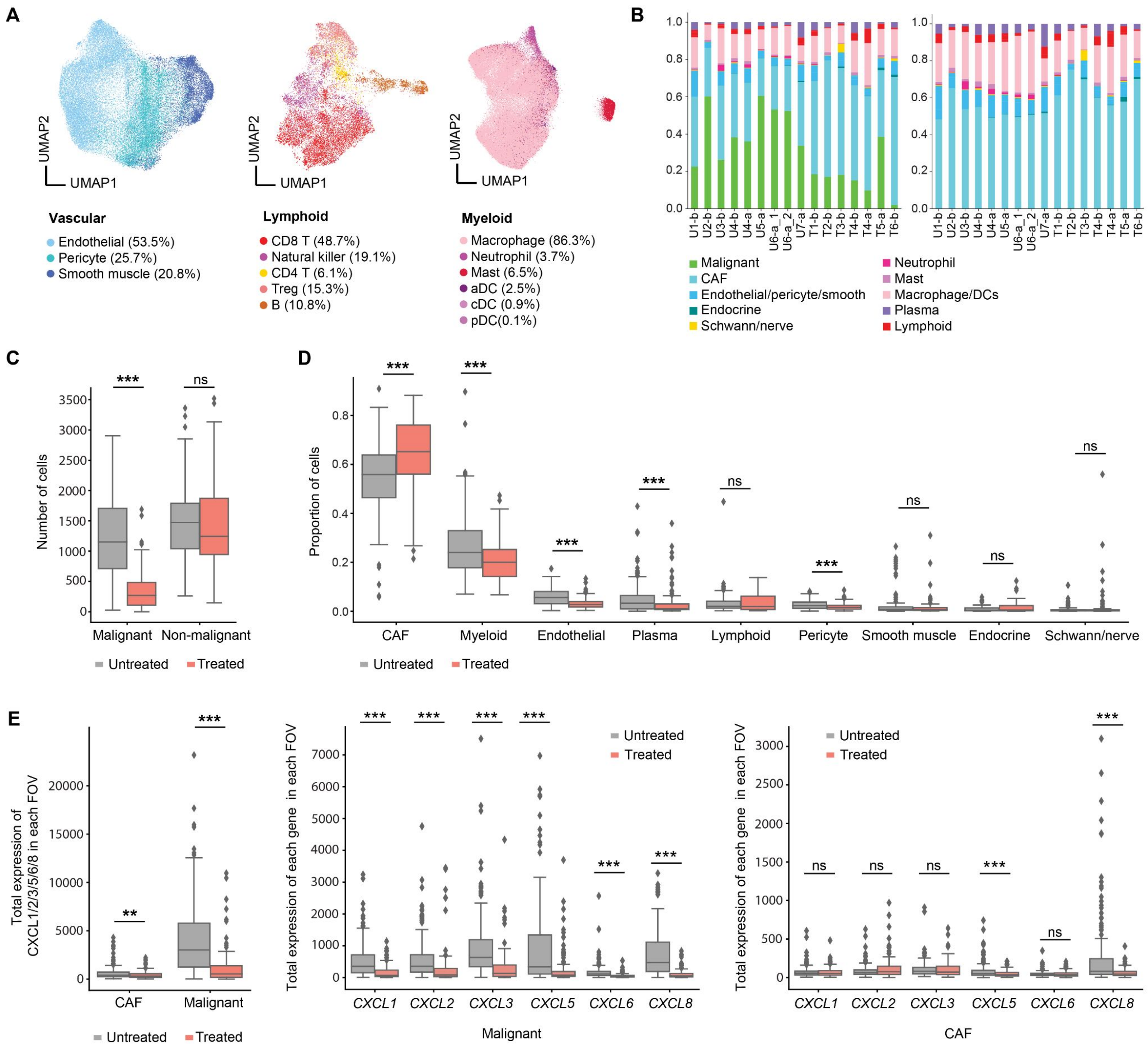

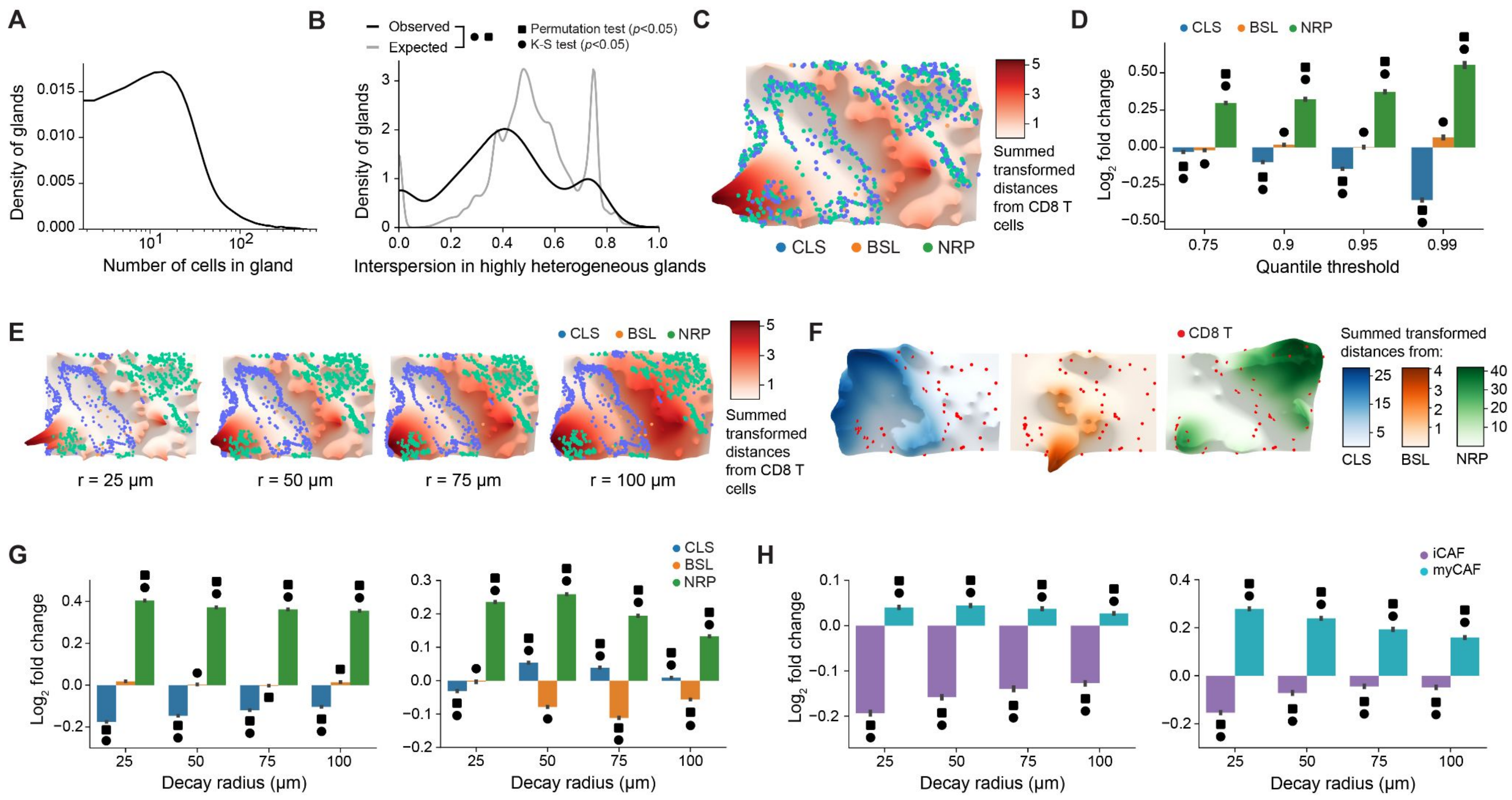

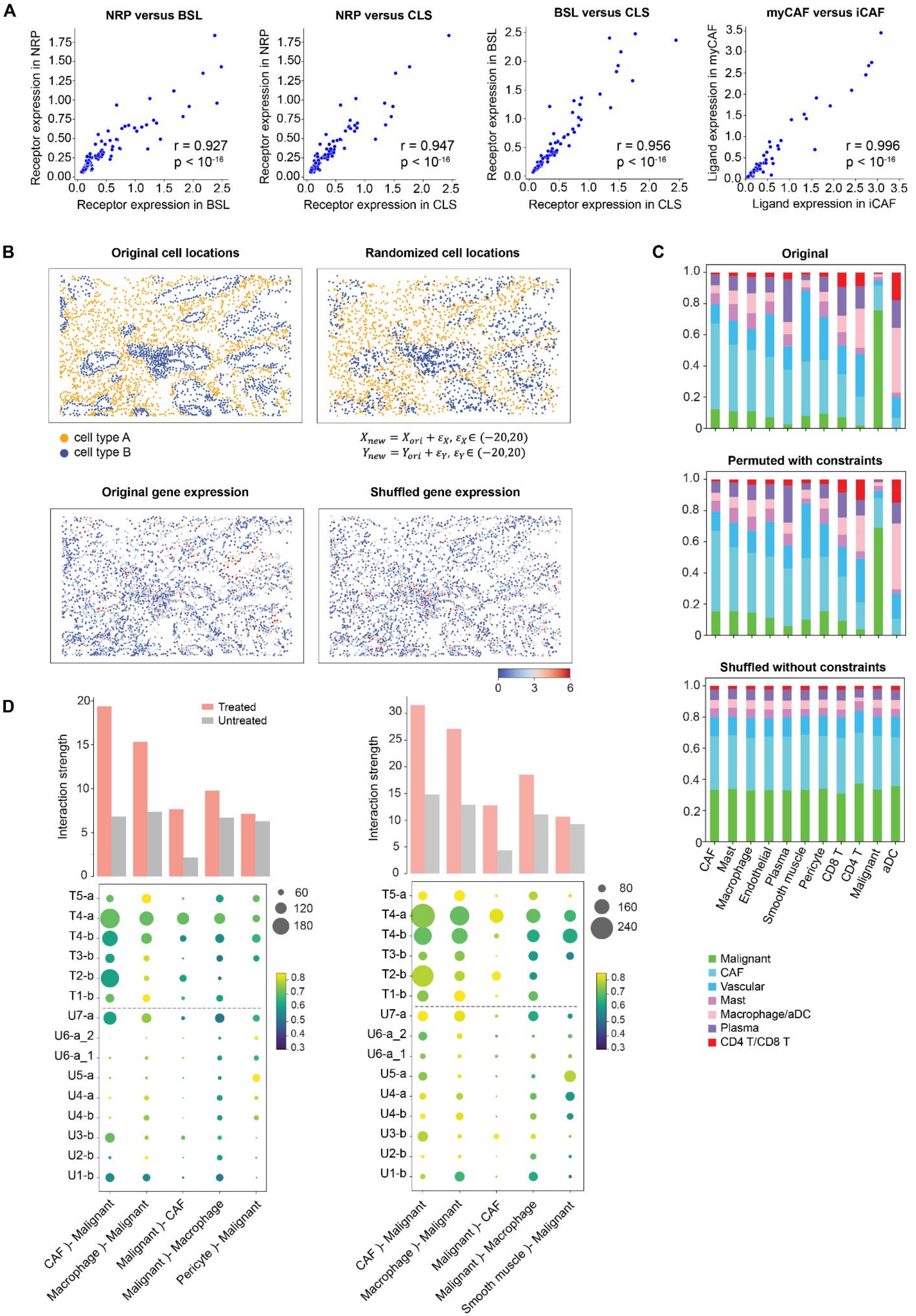

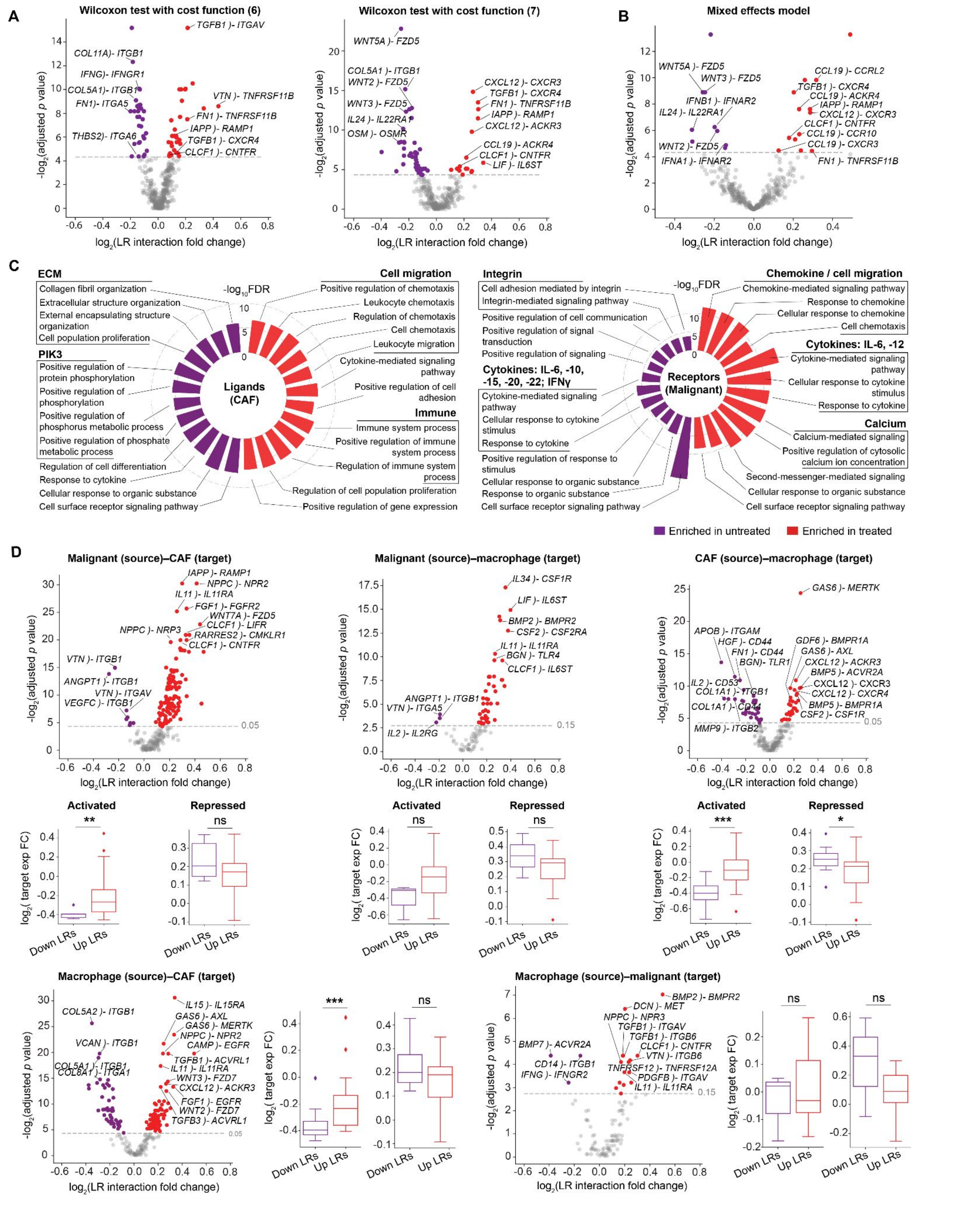

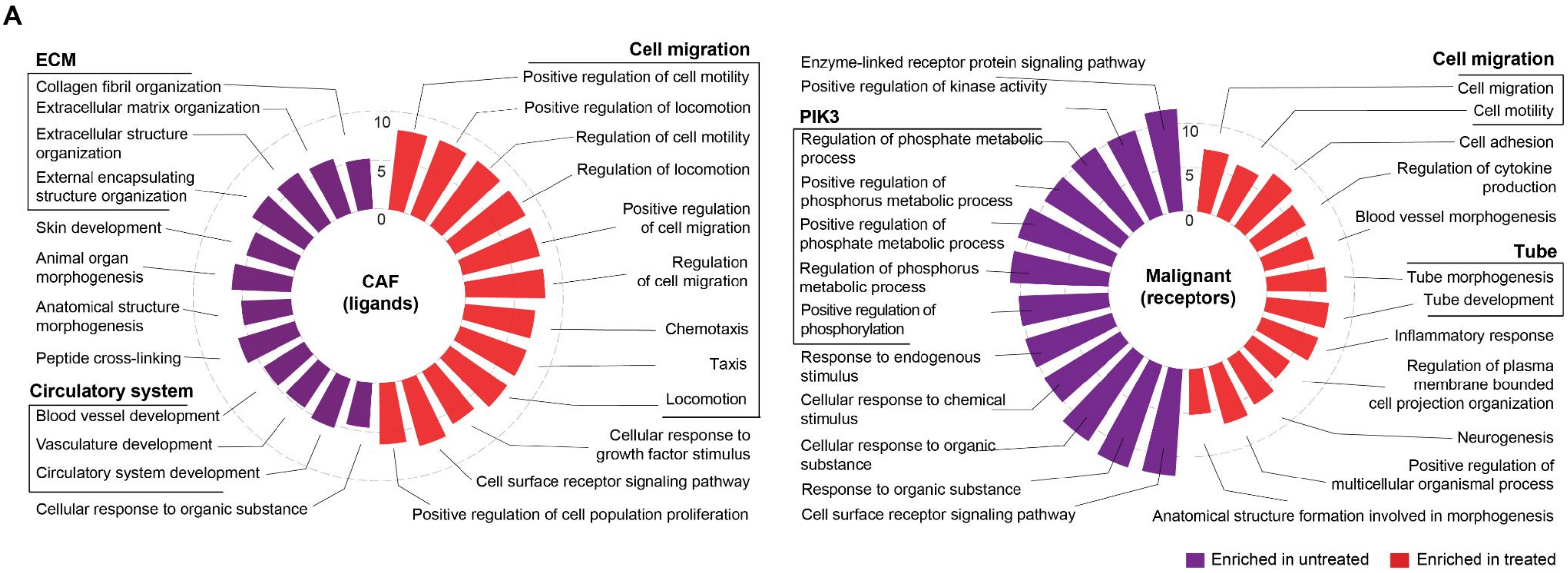

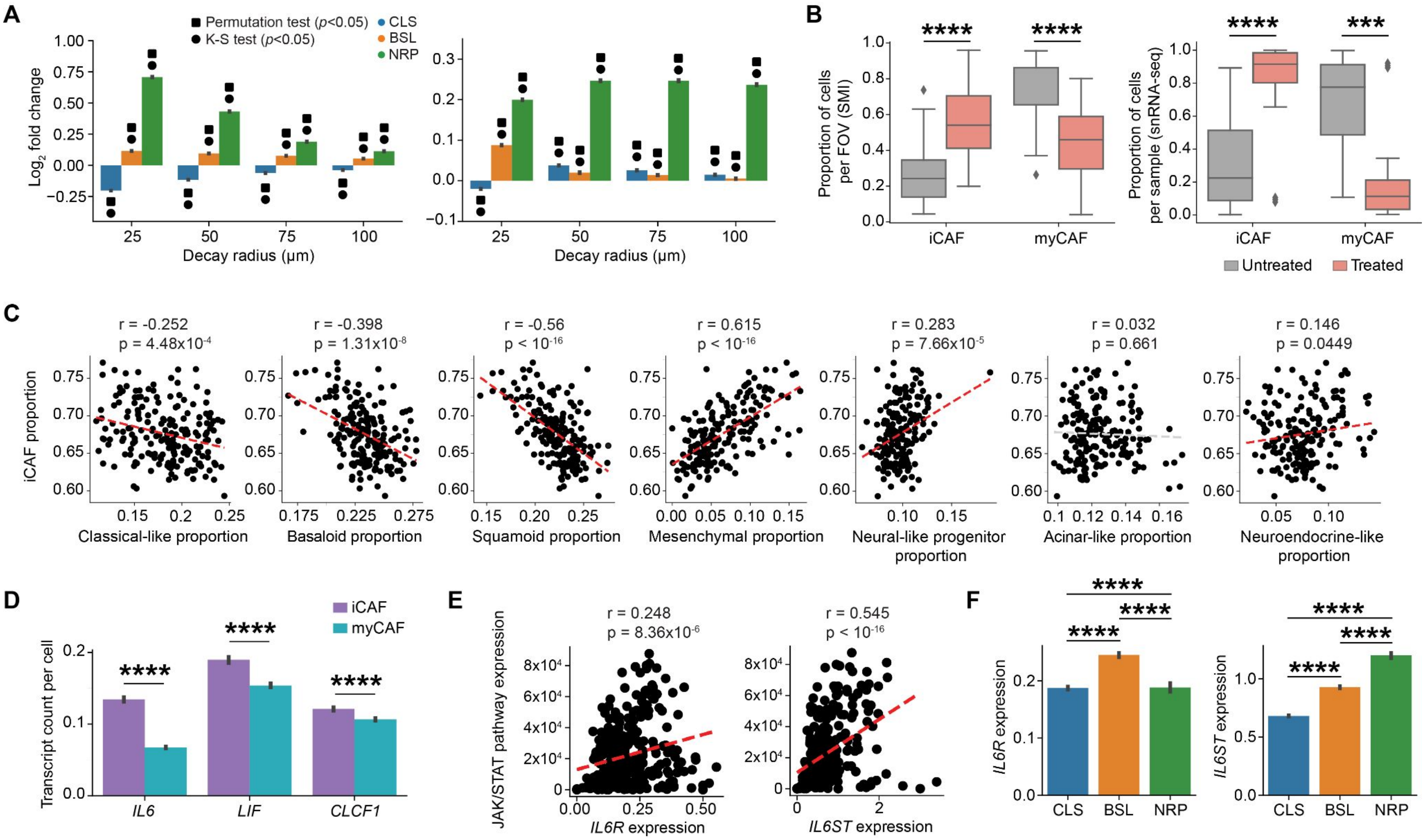
