## Supplementary material for "Therapy-associated remodeling of pancreatic cancer revealed by single-cell spatial transcriptomics and optimal transport analysis": Table S1

**Table S1.** Clinicopathological parameters and quality control metrics for each sample.

| Sample | Age (decade) | Sex | Stage/grade | Margin | Histology | Neoadjuvant | Status last FUP | PFS (d) | OS (d) | Neutrophil count (per µL) | Probe set | Matched snRNA-seq | Matched DSP WTA |
|---|---|---|---|---|---|---|---|---|---|---|---|---|---|
| U1-b | 60s | M | T3N1M0/g1 | R0 | | Untreated | DWD | 684 | 1417 | 4480 | Base | None | None |
| U2-b | 70s | M | T3N1M0/g2 | R0 | | Untreated | NED | 12 | 12 | 9140 | Base | None | None |
| U3-b | 60s | M | T3N0M0/gX | R0 | | Untreated | DWD | 104 | 2281 | 6910 | Base | None | None |
| U4-b | 70s | M | T2N1M0/g2 | R0 | | Untreated | MET | 35 | 607 | 3943 | Base | Present | Present |
| U4-a | 70s | M | T2N1M0/g2 | R0 | | Untreated | MET | 35 | 607 | 3943 | Augmented | Present | Present |
| U5-a | 80s | M | T3N2M0/g3 | R1 | AS | Untreated | DWD | -- | 171 | 820 | Augmented | Present | Present |
| U6-a_1 | 80s | M | T2N1M0/g2-3 | R1 | | Untreated | DWOD | 223 | 223 | 10307 | Augmented | Present | Present |
| U6-a_2 | 80s | M | T2N1M0/g2-3 | R1 | | Untreated | DWOD | 223 | 223 | 10307 | Augmented | Present | Present |
| U7-a | 70s | F | T2N0M0/g2-3 | R0 | | Untreated | NED | 112 | 112 | 3589 | Augmented | None | Present |
| T1-b | 60s | M | ypT2N2M0/g3 | R0 | AS | CRT | MET | 34 | 34 | 5747 | Base | Present | Present |
| T2-b | 70s | M | ypT3N0M0/g2 | R0 | | CRT | MET | 310 | 849 | 2273 | Base | Present | Present |
| T3-b | 70s | M | ypT3N0M0/g2 | R0 | | CRT | NED | 201 | 201 | 2928 | Base | Present | Present |
| T4-b | 70s | M | ypT2N0M0/g2 | R1 | | CRT | NED | 391 | 391 | 2840 | Base | Present | Present |
| T4-a | 70s | M | ypT2N0M0/g2 | R1 | | CRT | NED | 391 | 391 | 2840 | Augmented | Present | Present |
| T5-a | 70s | F | ypT1cN0M0/g2-3 | R0 | | CRT | DWD | 265 | 614 | 2345 | Augmented | None | Present |
| T6-b | 60s | M | ypT3N0M0/gX | R1 | | CRTL | NED | 1125 | 1125 | 1730 | Base | Present | Present |

**Table S1 (c*ont*).** Clinicopathological parameters and quality control metrics for each sample.

| Sample | Number of cells | Cells passed QC (%) | Number of transcripts | Number of FOVs | Transcripts per cell (mean) | Transcripts per cell (q10) | Transcripts per cell (q90) | Transcripts per cell per target (mean) | Negative probes per cell (mean) | Negative probes per cell per target (mean) |
|---|---|---|---|---|---|---|---|---|---|---|
| U1-b | 75640.0 | 81.8 | 17311721.0 | 24.0 | 161.2 | 42.0 | 328.0 | 0.168 | 0.863 | 0.0454 |
| U2-b | 60873.0 | 79.7 | 20087692.0 | 25.0 | 252.4 | 31.0 | 596.0 | 0.263 | 1.06 | 0.0559 |
| U3-b | 70451.0 | 86.5 | 24150265.0 | 24.0 | 242.9 | 48.0 | 492.0 | 0.253 | 0.600 | 0.0316 |
| U4-b | 67767.0 | 95.4 | 46046927.0 | 22.0 | 417.6 | 99.0 | 828.0 | 0.435 | 0.889 | 0.0467 |
| U4-a | 60821.0 | 95.5 | 35886399.0 | 20.0 | 526.5 | 109.0 | 1091.0 | 0.532 | 0.828 | 0.0436 |
| U5-a | 75975.0 | 98.4 | 65417212.0 | 24.0 | 786.0 | 188.0 | 1590.0 | 0.794 | 1.22 | 0.0642 |
| U6-a_1 | 83949.0 | 93.7 | 40675621.0 | 23.0 | 443.5 | 86.0 | 897.0 | 0.448 | 0.574 | 0.0302 |
| U6-a_2 | 69623.0 | 98.3 | 48820339.0 | 20.0 | 645.6 | 175.0 | 1242.0 | 0.652 | 1.034 | 0.0546 |
| U7-a | 93546.0 | 75.8 | 23270131.0 | 25.0 | 224.6 | 31.0 | 544.0 | 0.227 | 0.568 | 0.0299 |
| T1-b | 34239.0 | 91.6 | 22313009.0 | 22.0 | 324.2 | 69.0 | 647.0 | 0.338 | 0.866 | 0.0456 |
| T2-b | 50976.0 | 86.2 | 19664855.0 | 31.0 | 177.8 | 49.0 | 346.0 | 0.185 | 0.695 | 0.0366 |
| T3-b | 36480.0 | 92.7 | 22658662.0 | 21.0 | 305.8 | 74.0 | 613.0 | 0.319 | 0.775 | 0.0408 |
| T4-b | 46969.0 | 90.7 | 25789159.0 | 18.0 | 301.6 | 64.0 | 580.0 | 0.314 | 1.41 | 0.0744 |
| T4-a | 56312.0 | 98.3 | 33940791.0 | 21.0 | 490.0 | 148.0 | 906.0 | 0.495 | 0.959 | 0.0505 |
| T5-a | 46770.0 | 93.6 | 21532335.0 | 23.0 | 389.9 | 80.0 | 832.1 | 0.394 | 0.657 | 0.0346 |
| T6-b | 26211.0 | 79.5 | 10983912.0 | 16.0 | 189.8 | 23.0 | 391.0 | 0.198 | 1.11 | 0.0586 |
