## Supplementary material for "Therapy-associated remodeling of pancreatic cancer revealed by single-cell spatial transcriptomics and optimal transport analysis": Table S2

**Table S2.** Overlapping and unique ligand-receptor pairs inferred by using different databases.

| Ramilowski CellphoneDB intersection | Ramilowski CellchatDB intersection | CellphoneDB CellchatDB intersection | Ramilowski unique | CellphoneDB unique | CellchatDB unique |
|---|---|---|---|---|---|
| LRs enriched in treated samples | | | | | |
| CCL19_ACKR4 | CCL19_ACKR4 | CCL19_ACKR4 | VTN_TNFRSF11B | WNT3_WIF1 | COL9A1_ITGAV |
| LIF_IL6ST | LIF_IL6ST | LIF_IL6ST | FN1_TNFRSF11B | WNT7A_WIF1 | COL6A1_ITGB8 |
| CLCF1_CNTFR | CLCF1_CNTFR | CLCF1_CNTFR | CXCL12_CXCR3 | WNT7B_WIF1 | |
| IL23A_IL12RB1 | IL23A_IL12RB1 | IL23A_IL12RB1 | CSF2_CSF3R | WNT2_WIF1 | |
| CCL19_CCRL2 | VTN_ITGB5 | WNT7B_FZD4 | CCL19_CCR10 | WNT9A_FZD4 | |
| IAPP_RAMP1 | IL16_CD4 | | TGFB1_CXCR4 | PTGDS_PTGDR2 | |
| TGFB1_ITGAV | CXCL12_ACKR3 | | CCL19_CXCR3 | IGF1_ITGB4 | |
| | | | PDGFB_ITGAV | CXCL12_DPP4 | |
| | | | CXCL12_CD4 | | |
| | | | PF4_CXCR3 | | |
| | | | CCL2_ACKR4 | | |
| | | | IL16_CCR5 | | |
| LRs enriched in untreated samples | | | | | |
| OSM_OSMR | OSM_OSMR | OSM_OSMR | ANGPT1_ITGA5 | WNT11_FZD6 | ANGPT2_ITGB1 |
| WNT2_FZD5 | WNT2_FZD5 | WNT2_FZD5 | ANGPT1_ITGB1 | COL3A1_ITGA1 | TGFB2_ACVR1B |
| WNT3_FZD5 | WNT3_FZD5 | WNT3_FZD5 | CD14_ITGB1 | COL5A1_ITGA2 | WNT10B_FZD5 |
| WNT5A_FZD5 | WNT5A_FZD5 | WNT5A_FZD5 | COL1A1_ITGA5 | COL5A2_ITGA2 | WNT10B_FZD6 |
| WNT5A_FZD6 | WNT5A_FZD6 | WNT5A_FZD6 | COL3A1_DDR1 | COL11A1_ITGA1 | ANGPTL1_ITGA1 |
| IL10_IL10RB | IL10_IL10RB | IL10_IL10RB | COL4A5_CD47 | COL21A1_ITGA1 | ANGPTL1_ITGB1 |
| IL15_IL2RB | IL15_IL2RB | IL15_IL2RB | COL5A2_DDR1 | COL21A1_ITGB1 | |
| IL15_IL15RA | IL15_IL15RA | IL15_IL15RA | COL11A1_DDR1 | COL27A1_ITGA1 | |
| IL20_IL20RA | IL20_IL20RA | IL20_IL20RA | COL18A1_ITGA5 | COL27A1_ITGB1 | |
| IL20_IL22RA1 | IL20_IL22RA1 | IL20_IL22RA1 | CSF2_ITGB1 | COL8A1_ITGB1 | |
| IL24_IL22RA1 | IL24_IL22RA1 | IL24_IL22RA1 | GZMB_IGF2R | COL12A1_ITGB1 | |
| IFNA1_IFNAR2 | IFNA1_IFNAR2 | IFNA1_IFNAR2 | IL24_IL20RA | COL15A1_ITGB1 | |
| IFNB1_IFNAR2 | IFNB1_IFNAR2 | IFNB1_IFNAR2 | THBS2_ITGA6 | COL16A1_ITGB1 | |
| IFNG_IFNGR1 | IFNG_IFNGR1 | IFNG_IFNGR1 | VCAN_SELL | | |
| COL1A1_ITGA1 | COL1A1_ITGA1 | COL1A1_ITGA1 | VCAN_TLR2 | | |
| COL1A1_ITGB1 | COL1A1_ITGB1 | COL1A1_ITGB1 | VCAN_ITGB1 | | |
| COL1A2_ITGA1 | COL1A2_ITGA1 | COL1A2_ITGA1 | VEGFC_ITGB1 | | |
| COL1A2_ITGB1 | COL1A2_ITGB1 | COL1A2_ITGB1 | VTN_ITGA5 | | |
| COL6A3_ITGB1 | COL6A3_ITGB1 | COL6A3_ITGB1 | VTN_CD47 | | |
| FN1_ITGA5 | FN1_ITGA5 | FN1_ITGA5 | WNT5A_FZD8 | | |
| VTN_ITGB1 | VTN_ITGB1 | VTN_ITGB1 | | | |
| GDF9_BMPR2 | GDF9_BMPR2 | GDF9_BMPR2 | | | |
| WNT5A_RYK | NPPC_NPR2 | WNT7B_FZD6 | | | |
| IFNL3_IL10RB | THBS2_ITGB1 | WNT5B_FZD5 | | | |
| COL3A1_ITGB1 | THBS2_CD47 | WNT7B_FZD5 | | | |
| COL5A1_ITGA1 | | WNT9A_FZD5 | | | |
| COL5A1_ITGB1 | | WNT11_FZD5 | | | |
| COL5A2_ITGA1 | | COL9A2_ITGA1 | | | |
| COL5A2_ITGB1 | | COL9A2_ITGA2 | | | |
| COL11A1_ITGB1 | | COL9A1_ITGB1 | | | |
| COL18A1_ITGB1 | | COL9A2_ITGB1 | | | |
| | | COL9A3_ITGB1 | | | |
